# Pooled tagging and hydrophobic targeting of endogenous proteins for unbiased mapping of unfolded protein responses

**DOI:** 10.1101/2023.07.13.548611

**Authors:** Stephanie E. Sansbury, Yevgeniy V. Serebrenik, Tomer Lapidot, George M. Burslem, Ophir Shalem

**Affiliations:** Center for Cellular and Molecular Therapeutics, Children’s Hospital of Philadelphia, Philadelphia, PA 19104, USA; Department of Genetics, Perelman School of Medicine, University of Pennsylvania, Philadelphia, PA 19104, USA; Department of Biochemistry and Biophysics, Perelman School of Medicine, University of Pennsylvania, Philadelphia, PA 19104, USA; Department of Cancer Biology, Perelman School of Medicine, University of Pennsylvania, Philadelphia, PA 19104, USA; Epigenetics Institute, Perelman School of Medicine, University of Pennsylvania, Philadelphia, PA 19104, USA

**Author notes:** These authors contributed equally to this work (Stephanie E. Sansbury, Yevgeniy V. Serebrenik). Correspondence (Y.V.S.), (O.S.).

## Abstract

System-level understanding of proteome organization and function requires methods for direct visualization and manipulation of proteins at scale. We developed an approach enabled by high-throughput gene tagging for the generation and analysis of complex cell pools with endogenously tagged proteins. Proteins are tagged with HaloTag to enable visualization or direct perturbation. Fluorescent labeling followed by *in situ* sequencing and deep learning-based image analysis identifies the localization pattern of each tag, providing a bird’s-eye-view of cellular organization. Next, we use a hydrophobic HaloTag ligand to misfold tagged proteins, inducing spatially restricted proteotoxic stress that is read out by single cell RNA sequencing. By integrating optical and perturbation data, we map compartment-specific responses to protein misfolding, revealing inter-compartment organization and direct crosstalk, and assigning proteostasis functions to uncharacterized genes. Altogether, we present a powerful and efficient method for large-scale studies of proteome dynamics, function, and homeostasis.

## Introduction

Large scale imaging and gene perturbation technologies have transformed cell biology but are still very limited to nucleic acids. This is due to the ease of generating targeting reagents that are specific, using simple sequence complementarity, which is lacking from the protein space. For example, while advanced use of fluorescence *in situ* hybridization now enables tracking the subcellular localization of thousands of transcripts^1^, similar approaches for proteins are much more limited in multiplexing and scale^2,3^. A similar challenge exists when perturbing genes at scale: while the DNA and RNA space is easily accessible to perturbation reagents that are very specific, as in CRISPR and RNA interference^4–6^, proteins cannot be perturbed in living cells in a similar manner. The ability to target proteins at scale will enable studies into acute protein disruption, which is less prone to cellular adaptation or growth depletion when the targeted protein is lost with slow temporal dynamics^7^. In addition, direct protein targeting could induce many additional effects including stabilization, changes in subcellular localization, protein misfolding, and more^8^.

One widely used approach to both visualize and target proteins with high specificity is by generating fusions of the protein of interest with fluorescent domains, epitope tags, or degrons. Genome editing tools have simplified the generation of knock-in gene fusions, enabling studies of proteins in the endogenous regulatory and expression context^9^. These advancements have enabled the generation of large libraries of endogenous fluorescent fusions, demonstrating that they can provide unique and crucial insights into cellular organization and protein function^10^. Because these approaches still rely on the delivery of gene-specific reagents (e.g., homology dependent repair templates) in an arrayed format, they remain labor intensive and limited in scale. To address this, we set out to develop an approach that would enable performing such experiments in a pool by utilizing generic reagents^11^. Our method relies on the use of a synthetic exon inserted into introns in a homology-independent manner. This approach offers several advantages: reduced susceptibility to disruption by indels, a wide range of potential targeting sites, the ability to test multiple tag sites per gene due to the abundance of introns, and facile insertion of large fusion domains^11^. Indeed, this approach has been used successfully to isolate a large array of cell clones with endogenously tagged proteins from a tagging pool^12^.

Here, we develop a platform for Scalable POoled Targeting with a LIgandable Tag at Endogenous Sites (SPOTLITES). Using SPOTLITES, we recovered multiple nondisruptive tag sites for the vast majority of targeted genes, generating a cell pool with diverse localization patterns. To enable both protein visualization and perturbation, we use HaloTag as our donor domain. HaloTag covalently binds ligands with a chloroalkane linker that can be attached to a variety of small molecules for both imaging and direct protein targeting^8,13^. We utilized SPOTLITES capacity for high throughput imaging and protein perturbation to study the cellular response to localized unfolded protein stress in an unbiased manner. Cells have evolved intricate and complex molecular mechanisms for cellular proteostasis, a term referring to the maintenance of a healthy and functional proteome. Proteostasis involves tailored, compartment specific stress response pathways that are activated in the presence of protein folding stress. Some compartment-specific unfolded protein responses are well studied, such as those in the endoplasmic reticulum (ER)^14,15^ and mitochondria^16^, but many are poorly understood or completely unknown. Moreover, differences in proteostasis mechanisms within compartments are even more obscure despite the well-documented heterogeneity of most compartments in the cell^17–19^. A major difficulty in studying compartment-specific folding stress is a lack of means to specifically target perturbations. Often, pharmacological tools are used with nonspecific and/or toxic side-effects. Recently, HaloTag-localized hydrophobic tagging has emerged as a potent and highly specific way to induce spatially restricted protein misfolding, unraveling proteostasis mechanisms in the cytosol, ER, and even the Golgi apparatus^20–24^.

To induce unfolded protein stress in specific compartments and subcompartments throughout the cell, we treated an endogenously tagged cell pool with the hydrophobic HaloTag ligand HyT36^20^, followed by single cell RNA sequencing (scRNAseq) to measure the response to acute localized proteotoxic stress. The same cell library was also treated with a HaloTag TMR ligand for imaging, followed by *in situ* sequencing of sgRNA spacer sequences^25^ and computational clustering of protein localization patterns. By analysis of the response to protein misfolding of each individual localization cluster, we were able to identify compartment-specific transcriptional responses in an unbiased manner. This analysis revealed unique relationships and extensive crosstalk between certain cellular compartments. Reassuringly, we identify the well-characterized ER unfolded protein response in ER-localized tags, but also clearly in mitochondrial tags, highlighting the direct physical interaction between these two organelles. Many proteostasis-specific gene sets show unique response profiles across compartments including proteasomal and ribosomal genes. Lastly, we use our unique scRNAseq data set to assign a proteostasis function to uncharacterized stress responsive genes. We also provide a freely available website for the easy design of intron tagging libraries (www.pooledtagging.org). Altogether, we introduce a powerful new platform to measure and perturb endogenous proteome dynamics, opening the door for novel functional and imaging-based proteomics approaches.

## Results

### Pooled intron tagging enables the generation of complex cell libraries with endogenously tagged proteins spanning diverse localization patterns

We set out to test the feasibility of generating complex pooled libraries of cells with endogenously tagged proteins in a manner that is akin to typical low multiplicity of infection (MOI) pooled CRISPR screens. To achieve this goal, we adapted our previously published homology-independent intron tagging approach^11^ such that sgRNAs are delivered using low MOI lentiviral transduction followed by transfection of the following generic reagents: 1) a plasmid expressing Cas9, 2) a donor plasmid containing HaloTag flanked by constitutive splicing acceptor and donor sites, and 3) a plasmid expressing an sgRNA to linearize the donor plasmid in cells (Fig. 1A). We reasoned that by testing many possible tag sites for each protein, utilizing all introns, we would be able to identify non disruptive tagging locations for a majority of target genes. We decided to target a set of 265 genes that were manually curated for high expression and diverse, compartment-specific localization patterns (Fig. S1A). Our pooled sgRNA library was designed computationally to optimize sgRNA specificity and spacing across introns, and limit the number of sgRNAs per intron to 10 (Fig. S1B-C) resulting in a library of 21,345 sgRNAs. We created metrics to standardize this library based on gene coverage, sgRNA quality, and spacing within introns (Fig. S1D-H). sgRNAs were divided into three sublibraries based on coding frames (F0, F1, and F2) and matched with the corresponding donor to add the tag in frame. We also added 999 non-targeting control sgRNAs split proportionally across the sublibraries which served as negative controls for tagging.

**Figure 1.**
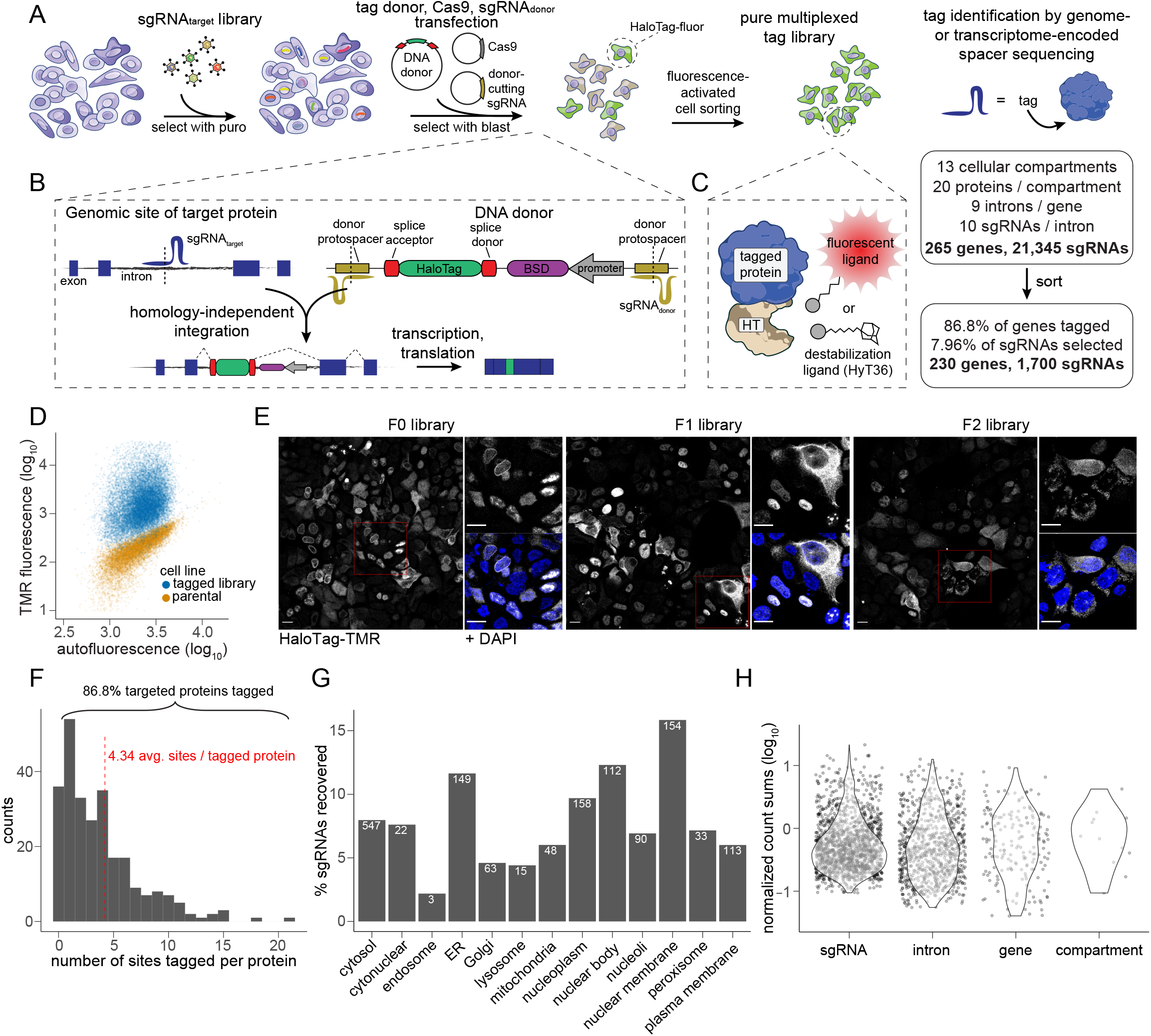
Pooled intron tagging enables the generation of complex cell libraries with endogenously tagged proteins spanning diverse localization patterns. A) Workflow for pooled tag cell library generation and analysis by sequencing. Right panel: designed vs sort-enriched library composition. B) Tagging is performed by homology-independent intron targeting, which entails transfection of generic reagents including a HaloTag donor encoding an independently-expressed selection marker. C) The FACS-purified library can be treated with various HaloTag ligands, including a fluorescent ligand for visualization or a hydrophobic tag (HyT36) for perturbation. D) Flow cytometry-based visualization of the F0 pooled tag library treated with HaloTag-TMR in comparison to the parental cell line. E) Confocal imaging of the three phase-specific pooled tag libraries treated with HaloTag-TMR to visualize individual tagged proteins. Magnified insets highlight details in the red boxes. Scale bars represent 10 μm. F) Histogram of the number of different sites (i.e., introns) tagged for each targeted protein in the total library. G) Percentage of sgRNAs recovered in the purified total library grouped by subcellular localization of the targeted protein. The numbers in each bar represent the total number of sgRNAs recovered. H) Normalized plots representing the number of sgRNA, intron, gene, or compartment elements recovered in the purified total tag library.

To test our approach, we performed our initial tagging in HAP1 cells. HAP1 cells were edited while in a haploid state to generate a high proportion of biallelic tags after reversion to a diploid state (Fig. S2A)^26,27^. Following low MOI transduction of the library, we delivered the generic components for tagging. The DNA donor, which was engineered in a minicircle plasmid, also contained a blasticidin resistance gene outside the splice sites with a dedicated promoter which was used to enrich cells with successful DNA integration events^11^ (Fig. 1B). Since cells were tagged with HaloTag, we could use a variety of HaloTag ligands to label the library (Fig. 1C). We used a fluorescent HaloTag ligand at a low concentration to sort cells and enrich for expressed and stable protein fusions (Fig. S3A-B, Fig. 1D). This also allowed us to track the purity of the library over multiple selection and sorting steps (Fig. S2B). Confocal microscopy visualization of the purified library revealed a wide variety of localization patterns corresponding to the targeted cell compartments (Fig. 1E). We further analyzed the cell library by bulk amplicon sequencing of the genomically integrated sgRNA cassettes in each cell, revealing that while ∼8% of designed sgRNAs led to tagging, ∼87% of all genes were tagged at least once (Fig. 1A, right panel). Protein fusions were recovered at an average of ∼4.3 different sites per gene (Fig. 1F), and there did not appear to be any major biases in tagging efficiency by compartment (Fig. 1G). Tagging efficiency can vary by orders of magnitude across target sites^11^, so we expected the resulting library to have a skewed representation of sgRNAs. Indeed, we find high skew at the level of sgRNAs and introns which is reduced at the level of the gene and compartment (Fig. 1H). Analysis of non-targeting control sgRNAs revealed a substantial depletion after sorting, consistent with our expectation that they would not lead to productive tagging (Fig. S2C).

We also tested pooled tagging in HEK 293 (Fig. S3) and HeLa cells (Fig. S7). After selection and sorting, both lines demonstrated diverse localization patterns of tags by microscopy (Fig. S3C, Fig. S7A). Bulk sequencing of HEK 293 cells at multiple steps during library generation revealed that most of the sgRNA skew is established during genomic integration followed by cell sorting (Fig. S3D). After sorting, tag libraries remain stable in composition for at least two weeks, permitting long-term experiments (Fig. S3E). We picked several sgRNAs for individual validation to confirm their quality and consistency of tagging. All produced specific and largely consistent localization patterns, even in polyclonal populations, as observed previously (Fig. S3F)^11^. In sum, we detail a quick and low-cost method for pooled endogenous protein tagging with high gene-level success and diversity.

### Exhaustive intron tagging selects for non-disruptive protein fusions

Our experimental procedure for the generation of pooled endogenously tagged libraries starts from a very large sgRNA library to produce a final library that contains a small subset of the designed sgRNAs. We next asked to what extent the selected tagging locations are non-disruptive to the function of the tagged protein, and if by analysis of the features of the successful sgRNAs, we can identify rules to guide the design of future libraries. To test this, we asked if growth essential genes can be tagged without leading to cell depletion, thus suggesting that growth-essential function is preserved. We tested this empirically by performing a CRISPR knockout growth screen on largely the same set of genes that we used for tagging (Fig. 2A). As expected, cells in which we knocked out essential genes rapidly depleted, while cells with sgRNAs against non-essential genes or non-targeting controls remained abundant (Fig. 2B). Comparing the extent of depletion of cells that are knockouts for an essential gene to that of cells with tags in the same essential gene revealed no significant correlation (Fig. 2C), suggesting that protein essentiality is not a major determinant for tagging, and moreover, tagging internally can be used while maintaining essential protein function (Fig. 2C). Similarly, we observed that essential genes, as annotated by DepMap^28^, tagged at single alleles in HEK 293 cells maintained stability for at least two weeks (Fig. S3E).

**Figure 2.**
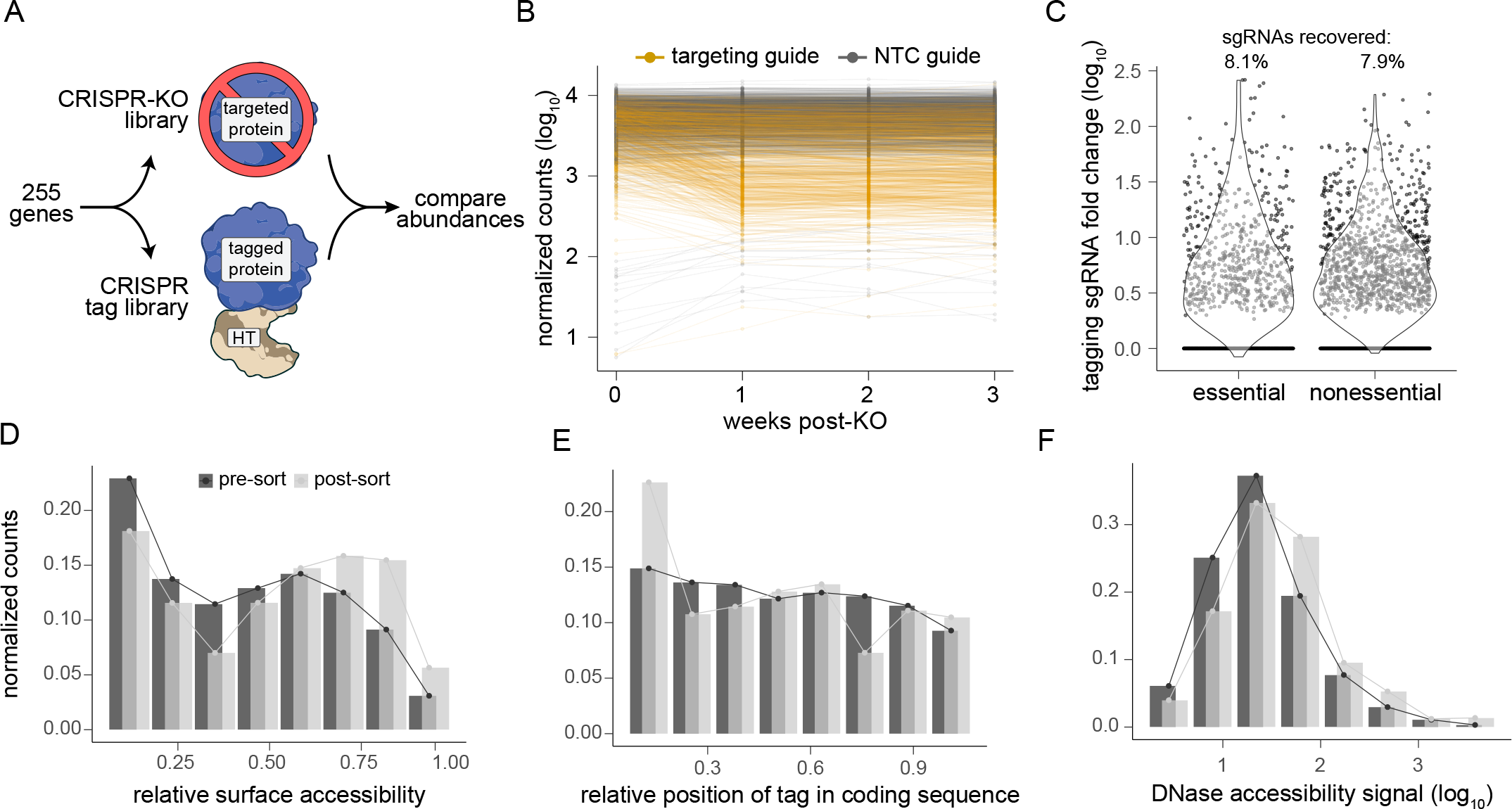
Exhaustive intron tagging selects for non-disruptive protein fusions. A) Diagram of experiment outline comparing gene tagging to gene knockout (KO). B) KO sgRNA abundance over time. C) Tagging sgRNA fold changes grouped by whether the targeted gene was determined to be essential or not. Violin plots represent the recovered sgRNAs. D-F) Abundance of sgRNAs targeting sites of various D) protein relative surface accessibilities, E) relative positions within the coding sequence, or F) genomic DNase accessibility, pre- and post-sorting.

We next asked which protein or genomic features are predictive of successful tagging. Exploring physical protein features, we find that tagging is enriched at surface-accessible residues compared to internally buried ones which we expect to be more likely to lead to destabilization and degradation of the fusion protein (Fig. 2D). Similarly, tagging is enriched within disordered protein domains where important structures are less likely to be disrupted (Fig. S4A-B). We also see that smaller proteins are easier to tag, potentially reflecting that fused HaloTag is more likely to fold and bind a fluorescent tag with less steric obstruction (Fig. S4C). Within the coding sequence, we see a strong preference for tagging near the protein N-terminus (Fig. 2E). This is most likely a reflection of the epigenetic state of the tagging site which is more open at transcription start sites and thus more likely to be successfully edited there^29,30^. In support of this observation, we also find enriched tagging at genomic sites with higher DNase accessibility (Fig. 2F), and at sites with stronger H3K4Me3 and H3K27Ac signals (Fig. S4D-E) which are markers of open chromatin^31,32^. Bulk RNA sequencing revealed that transcription is positively correlated with successful tagging, further emphasizing the link to epigenetic state (Fig. S4F-G). These results track closely with recent findings that chromatin accessibility has a significant impact on Cas9-associated editing activity^33,34^.

We found no obvious effect of the relative position of a tag site within an intron on tagging success (Fig. S4H-K). Considering the contribution of predicted sgRNA score^35^ to tagging success, we see only slight differences between the pre- and post-sort tag populations. We detect a slight preference for sgRNAs with higher efficiency scores (Fig. S4L) and intermediate specificity scores (Fig. S4M). Notably, our library utilized sgRNAs with higher efficiency and specificity scores (Fig. S1E), likely explaining why we don’t see a stronger influence of sgRNA scores on tagging. These analyses thus reveal relevant genomic-, protein-, and sgRNA-level features for tagging success which can be employed to design functional and high-efficiency *de novo* tag libraries in the future, in addition to the incorporation of empirically successful tagging sgRNAs which we hope to accumulate over time.

### High throughput imaging and in situ sequencing of the sgRNA in each cell enables systematic evaluation of localization patterns

To analyze tagged protein localization at the level of individual proteins and cells, we performed automated high throughput image acquisition followed by *in situ* sgRNA sequencing to identify the tagged protein in each cell (Fig. 3A). Tagged cells stained with nuclear and cytoplasmic dyes were imaged at 40X magnification using a motorized programmable microscope stage. Next, *in situ* sequencing was performed following a protocol based on pooled imaging CRISPR screens^36^. Briefly, the sgRNA sequence is reverse transcribed in cells followed by padlock capture and rolling circle amplification to generate localized amplified DNA of the spacer sequence. Sequencing by synthesis of the spacer is performed with consecutive automated imaging cycles (Fig. 3B) at 10X magnification^37^. We then filtered cells based on imaging and *in situ* sequencing quality parameters and randomly arranged a small number of individual cells for each sgRNA into “cell albums” (Fig. 3C). This analysis revealed the ability to tag proteins with distinct localizations including throughout subcompartments of the cytosol and nucleus, within mitochondria, and across the secretory pathway. sgRNA abundance from *in situ* sequencing was in good agreement with abundance from bulk amplicon sequencing (Fig. S5A).

**Figure 3.**
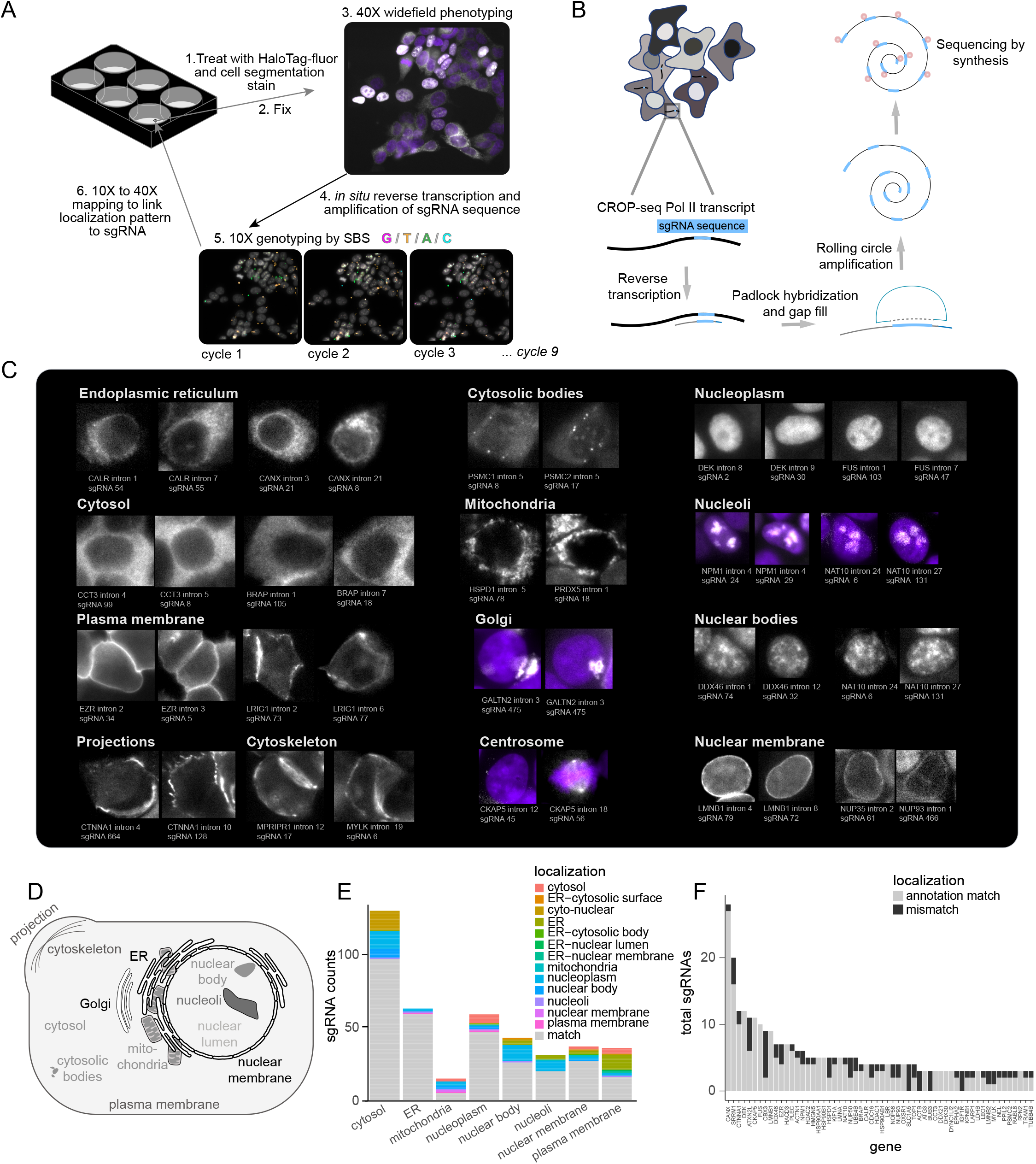
High throughput imaging and *in situ* sequencing of the sgRNA in each cell enables systematic evaluation of localization patterns. A) Workflow for live-cell HaloTag-fluor staining, fixation, phenotyping, and pooled sequencing. B) Detailed schematic of *in situ* reactions to generate fluorescent sgRNA sequences for imaging-based readout. C) Cell images and corresponding tagged protein identities for compartment-specific genes identified by optical pooled sequencing. D) Cartoon summary of the compartment-specific localization patterns represented in the pooled tagged cell library. E) Quantification of the proportion of compartment-specific sgRNAs that exhibit localization patterns consistent with literature-sourced annotation. sgRNAs that match the expected pattern are represented in gray with additional colors representing non-matching localization patterns. F) Counts of sgRNAs that match or do not match the annotated compartment on a per-gene level. Genes represented by at least three abundant sgRNAs are plotted.

Our original library was designed using proteins with established localization patterns such that we could explore the agreement between the expected and observed pattern for each tag. We grouped sgRNAs based on expected localization patterns clearly distinguished by imaging and examined whether the observed patterns matched. In most cases, the expected localization matched the observed one (Fig. 3E), yet the number of mismatched localization patterns was not negligible, suggesting that rigorous evaluation of localization patterns is important for the generation of high quality libraries. The proportion of mismatched sgRNAs per localization pattern did not vary dramatically between compartments, indicating that the compartment of the tagged protein is not an important determinant of tagging success (Fig. 3E). However, certain compartments were biased in the localization patterns of mismatches towards adjacent organelles; for example, mismatched nuclear body tags tended to look nucleoplasmic and mismatched plasma membrane tags tended to look ER-like (Fig. 3E). Analysis of sgRNA localization for each gene revealed that the majority of sgRNAs per gene resulted in appropriate localization (Fig. 3F), demonstrating that indeed, using our exhaustive intron tagging approach, we can identify multiple viable tagging sites for most proteins tested.

### Analysis of a compact tag library for exploring proteotoxic responses across cellular compartments

Our ability to reliably localize HaloTag to various cell compartments in a pool allowed us to use ligand-induced HaloTag misfolding to systematically assess how the cell responds to spatially-restricted proteotoxic stress. To do so, we combined pooled imaging of a high quality compact library with single cell RNA sequencing (scRNAseq) which reports on both transcriptional responses as well as the identity of the tag in each cell (Fig. 4A). To enable high scRNAseq depth and cell coverage for each tag with current reagents, we used sgRNAs that produced high-confidence tag localization patterns to construct a balanced and scaled down pooled library targeting 11 compartments with ∼3 genes each and ∼3 tag sites per gene (Table S1). This library is similar in compartment diversity to our previous library but uses only 146 instead of 22,344 sgRNAs. Tagging was again performed in haploid HAP1 cells (Fig. S6A) followed by purification by cell sorting (Fig. S6B). We have also used the same compact intron tagging sgRNA pool to construct a tagging cell library in HeLa cells (Fig. S7A). Comparison of the tag abundance between the cell lines demonstrated high correlation suggesting that tagging efficiency is highly reproducible, a fact that could be used for the design of future libraries with a more balanced tag representation (Fig. S7B).

**Figure 4.**
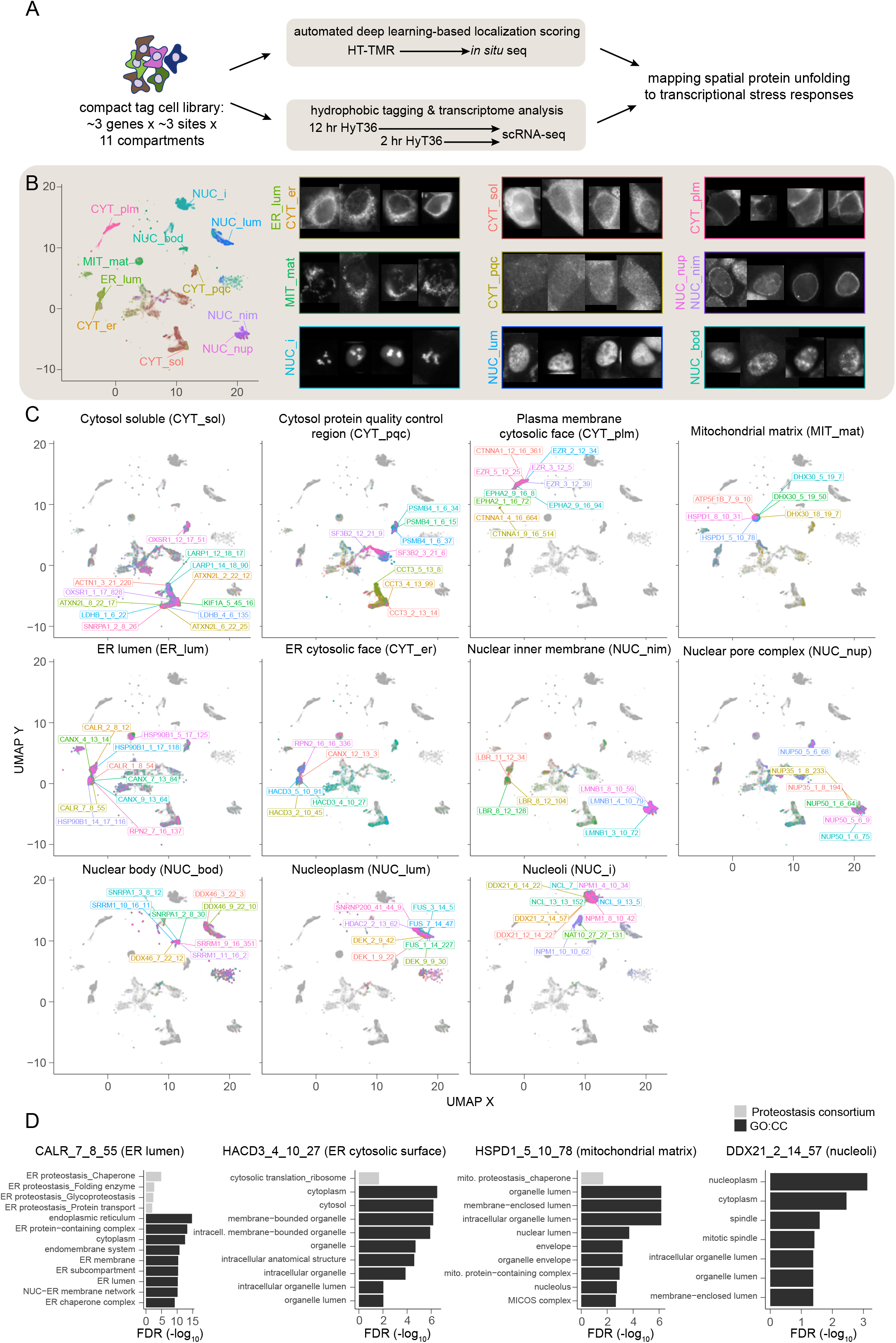
Analysis of a compact tag library for exploring proteotoxic responses across cellular compartments. A) Workflow of approach to achieve systematic and precise compartment-specific proteotoxic stress, performed on a compact and diverse endogenous tag library. The library is labeled with HaloTag-TMR (HT-TMR), analyzed by imaging and optical sequencing, and localization patterns are discerned by automated image analysis. In parallel, the library is labeled with the HaloTag hydrophobic tag, HyT36, to induce pooled protein misfolding, followed by analysis by single cell RNA sequencing (scRNAseq). Transcriptional responses are subsequently integrated with spatial information. B) Imaged cells are clustered by localization patterns using features extracted from a variational autoencoder and embedded onto a UMAP. Clusters represent distinct subcellular compartments depicted by cells sampled from the indicated cluster. C) Representation of tag groups and the associated sgRNAs, each mapped onto the UMAP and labeled at the mode of sgRNA density. sgRNA naming structure follows: *gene _ targeted intron # _ total # of introns _ sgRNA #*. D) Functional enrichment of GO:CC and Proteostasis Consortium terms in the ∼100 most upregulated genes per sgRNA in response to induced destabilization of the tagged proteins.

Following HAP1 library construction, we confirmed the localization pattern of each tag by high throughput imaging, *in situ* sequencing, and unbiased clustering (Fig. 4B-C, Fig. S8). We sampled a constant number of cells per sgRNA for deep learning-based localization analysis. Some filtering was applied to mitigate genotyping errors based on consensus localization patterns (STAR Methods; Supplementary Text). The Cytoself VAE^38^ was trained on selected cells and used to infer the global representation latent space parameters for every segmented cell image. The latent space was then embedded onto a two dimensional UMAP^39^, and clustered using HDBSCAN40 (STAR Methods; Supplementary Text). Each targeted compartment was represented by a unique cluster with consistent localization patterns (Fig. 4B). Compartments and subcompartments, together referred to as “tag groups,” consisted of sgRNAs determined by their empirical localization patterns which demonstrated high consistency and uniformity (Fig. 4C). For example, 11 of 12 tags targeting soluble cytosolic proteins centered at a single cluster with cytosolic localization and noticeable nuclear exclusion, while 1 tag centered at a related cytosolic cluster with weaker nuclear exclusion (Fig. 4C, panel 1). The related cluster is associated with the cytosolic protein quality control region tag group (CYT_pqc), which is composed of cytosolic proteins involved in protein quality control and that exhibit a proteasome gene-rich transcriptional response to misfolding (Fig. S11). Some compartments with overlapping localization patterns were intersected with literature annotation for more precise grouping; for example, nuclear membrane tags were separated into nuclear pore and nuclear inner membrane tag groups. All tags matched the literature-expected localizations with one exception, SF3B2, which exhibited a cytosolic pattern despite being annotated as nuclear (Fig. S8). Ultimately, SF3B2 tags were grouped with proteasome tags due to their transcriptional response to misfolding (Fig. S11). Occasional sgRNAs had patterns that split evenly between clusters, such as sgRNAs tagging LBR (lamin-B receptor). Closer examination of the localization patterns revealed an intermediate localization pattern which had a strong nuclear membrane pattern that extended partially into the ER, consistent with the clustering result (Fig. S8).

To induce HaloTag misfolding we utilized the hydrophobic tag HyT36 which is membrane permeable and has been used previously to study localized proteotoxic stress20–24. We optimized HyT36 treatment conditions using clonal HEK293 cell lines tagged via validated sgRNAs with various localization patterns (Fig. S3F). For tested clones, we targeted a HyT36 concentration that would cause subtle but noticeable protein depletion at 24 hours but minimal depletion before 10 hours (Fig. S9A-B). Similarly, to avoid off-target effects, we focused on concentrations that had minimal toxicity within the first 24 hours of treatment (Fig. S9C). We decided to treat cells with 25 μM HyT36 for 2 and 12 hours in order to capture the breadth of acute responses to misfolding stress across the different cell compartments. Prior to destabilization by HyT36 treatment, the three frame-specific tag sublibraries were pooled together at a ratio to minimize sgRNA count variance (Fig. S10A). We targeted 120,000 cells for 10x Genomics scRNAseq per treatment to guarantee at least 200x cell coverage for 56 tags or at least 50x cell coverage for 96 tags (Fig. S10B). Controls were sequenced at a lower cell coverage and included DMSO-treated tagged cell pools and a HyT36-treated untagged cell pool, which allowed us to rule out any adverse effects of DMSO or HyT36 treatment, respectively. An additional untagged cell pool was spiked into all samples after treatment but before single cell processing to serve as an internal control between GEM wells (STAR Methods). The read processing workflow and post-filter sample statistics are described in Fig. S10C. After normalization, all GEM wells for each treatment clustered closely together in a principal component analysis plot (Fig. S10D).

To determine differentially-expressed genes associated with the misfolding of each tag, cells were averaged per sgRNA and gene fold changes were calculated relative to a pseudo-random subset of the cell population (STAR Methods) (Fig. S10E). Of all controls available, this approach best accounted for non-specific effects of drug treatment and sample batch processing. To test the efficacy and specificity of our approach in a pathway-agnostic way, we examined the enrichment of GO:CC terms41,42 in the response associated with select individually tagged proteins (Fig. 4D). Reassuringly, HaloTag localized to the ER lumen by fusion to CALR led to enrichment of ER-associated GO:CC terms as well as ER proteostasis terms curated by the Proteostasis Consortium43,44 (Fig. 4D, panel 1). Conversely, HaloTag localized to the cytosolic surface of the ER did not lead to enrichment of ER terms but rather to cytosolic terms (Fig. 4D, panel 2) indicating that HaloTag produces proteotoxic stress that is spatially precise. HaloTag misfolded in the mitochondrial matrix induced upregulation of mitochondrial genes as well as mitochondrial chaperones (Fig. 4D, panel 3). Finally, HaloTag misfolded at the nucleolus led to the upregulation of nucleus-associated genes but interestingly no known proteostasis terms (Fig. 4D, panel 4). Similarly, systematic evaluation of the top upregulated genes for each sgRNA and their associated GO:CC terms revealed that protein misfolding often led to upregulation of genes associated with the stressed compartment at both 2 and 12 hours of treatment (Fig. S11A-B). We conclude that our integrated tag localization map and scRNAseq dataset provides a valuable resource to explore and discover new mechanisms within compartment- and subcompartment-specific transcriptional responses to proteotoxic stress.

### Analysis of transcriptional responses by localization clustering reveals signatures associated with spatially restricted proteotoxic stress

To characterize the responses to protein folding stress elicited by misfolding in different cell compartments, we analyzed the combined transcriptional response of each tag group. We first explored compartment-specific transcriptional responses by determining all genes that were significantly differentially expressed only in a single tag group (Fig. S12, Table 2). ER-localized tags functioned partly as positive controls since the ER UPR is well characterized15. For compartments with less characterized proteostasis pathways, we performed functional enrichment analysis of differentially expressed gene sets using broad annotations including GO:CC and GO:BP. Protein misfolding in the ER led to strong upregulation of genes associated with the ER and ER stress such as HSPA5 and several protein disulfide isomerases (Fig. S12A, Table 2). Similarly, protein misfolding in other compartments led to differential expression of compartment-specific genes, suggesting distinct and relevant proteostasis responses. For example, mitochondria-associated gene terms were enriched in response to protein misfolding in the mitochondria, and ncRNA metabolism was enriched in response to misfolding in nucleoli, consistent with observations from individual sgRNAs (Fig. 4D, Fig. S11). These gene sets thus provide insights into potentially novel compartment-specific unfolded protein responses.

To evaluate broad compartment similarity and relationships, we clustered tag groups by pairwise correlation of fold changes (Fig. 5A). We used a scale-invariant metric for comparison to minimize tag-specific differences in gene fold change magnitude. Furthermore, we focused on the top ∼3500 most significant differentially expressed genes (DEGs) for analysis (STAR Methods). Noticeably, we saw that each tag group was most closely related to itself at the alternate timepoint, suggesting a broad consistency to the transcriptional responses over the course of 10 hours. It was also apparent that tag groups separated into three “metagroups”, where tagged proteins associated with the ER lumen, mitochondrial matrix, and CYT_pqc formed the so-called “metagroup 1”, nuclear bodies and nucleoli formed “metagroup 2”, and the remaining nuclear and cytosolic tag groups formed “metagroup 3”. To better understand which functional gene categories are responsible for this clustering, we compared the two extreme metagroups, 1 and 3, in terms of their proteostasis-specific transcriptional responses. Proteostasis-specific genes, defined by the Proteostasis Consortium, were grouped into gene sets by their Branch and Class designations and intersected with the significant DEGs43,44. Most proteostasis genes were upregulated in metagroup 1 and downregulated in metagroup 3 (Fig. S13A). The gene set-specific significance of this difference was designated the “metagroup difference score” (Fig. 5B). We also explored which proteostasis gene sets demonstrated a tag group-specific response by comparing the maximum versus the absolute mean gene set-specific response across all tag groups (Fig. S13B). This comparison is expected to highlight gene sets that show a strong response which is confined to one tag group. Indeed, we find several ER-related gene sets to be the most specific, possibly reflecting that the ER has one of the most thoroughly detailed stress responses.

**Figure 5.**
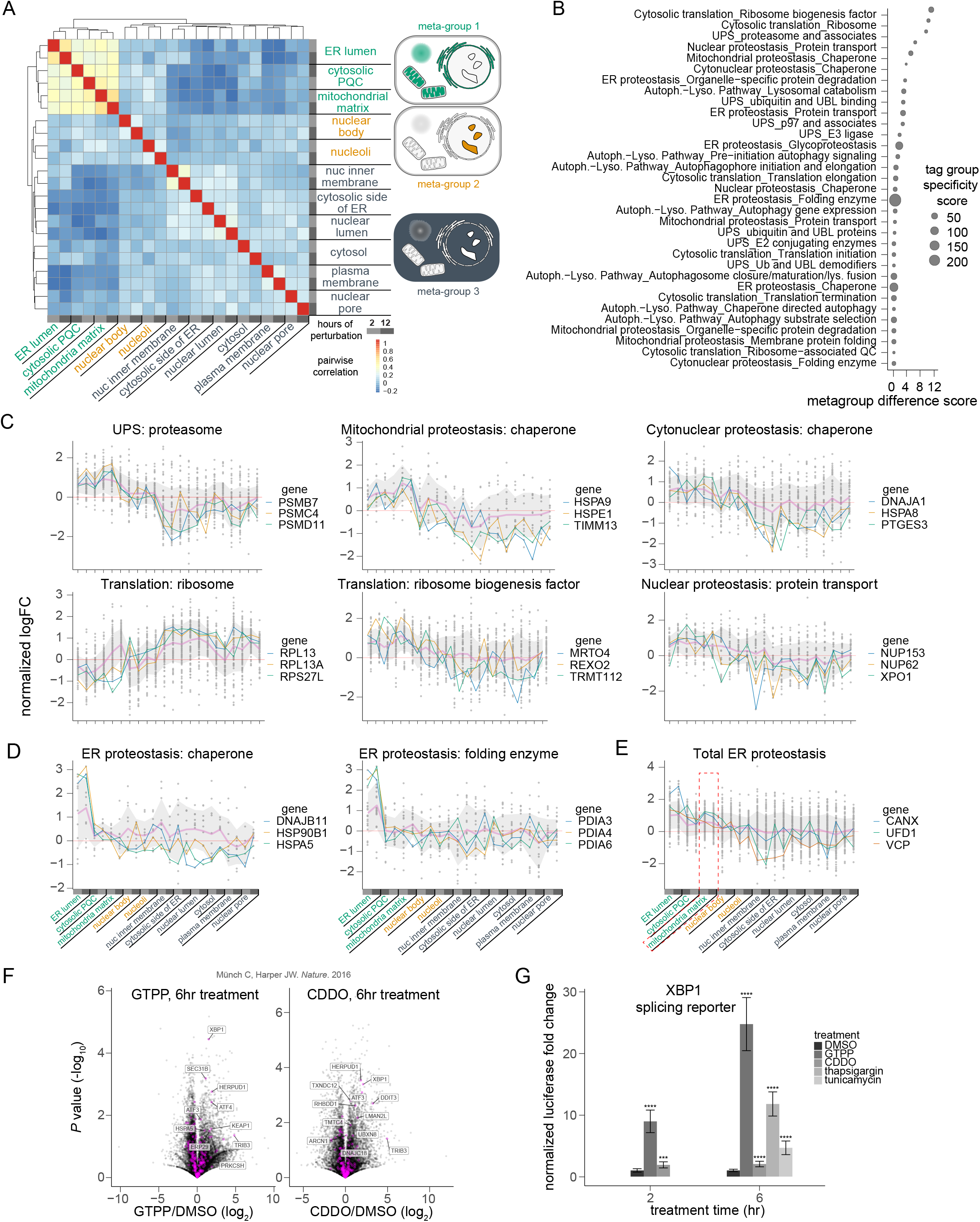
Analysis of transcriptional responses by localization clustering reveals signatures associated with spatially restricted proteotoxic stress. A) Left: heatmap of pairwise correlation values between tag groups at 2 and 12 hours of HyT36 treatment. Right: cell diagram depicting the different tag groups colored by metagroup which in turn is determined by hierarchical clustering. B) Proteostasis-associated gene sets ranked by metagroup difference and sized according to tag group specificity. C-E) Normalized log fold changes of proteostasis gene sets across tag groups ordered by hierarchical clustering from A. Points represent each gene in the labeled gene set, the thick pink line represents the mean of points, and the gray ribbon represents one standard deviation from the mean. For each plot, three genes with large differential expression between metagroups 1 and 3 are displayed. Six gene sets with the largest metagroup 1 and 3 differential are represented in C, and two gene sets with the largest tag group specificity are represented in D. All ER proteostasis-associated genes are depicted in E. F) Volcano plots of RNA sequencing data from the indicated study in human cells, with ER proteostasis-associated genes colored in magenta. G) XBP1 splicing-based luciferase reporter signal in cells with indicated treatments and normalized to cell viability. Asterisks represent corrected *P* values (**** < 0.0001, *** < 0.001).

We next explored the most strongly affected proteostasis gene sets in detail by visualizing the normalized fold change for each gene of a gene set across all tag groups (Fig. 5C-E). Most proteostasis gene sets, including proteasome subunits, various groups of chaperones, and ribosome biogenesis factors, were strongly upregulated for each tag group of metagroup 1, had low to intermediate upregulation in metagroup 2, and were downregulated in metagroup 3 (Fig. 5C). An inverse pattern was seen for ribosome subunits, which were downregulated in metagroup 1 and upregulated in metagroup 3 (Fig. 5C). Some gene sets showed more complex regulation, with subsets of genes responding in opposite directions (Fig. S14). Within autophagy, for example, macroautophagy genes demonstrated a positive correlation with proteostasis genes while mitophagy genes showed an inverse relationship (Fig. S14). Particularly for metagroup 1, all groups including the ER lumen, mitochondrial matrix, and CYT_pqc recapitulated responses well established in the literature. For example, ER proteostasis factors were most strongly upregulated in response to ER perturbation, mitochondrial chaperones in response to mitochondrial perturbation, and proteasome genes in response to perturbation of the CYT_pqc45.

Furthermore, across metagroup 1, we saw a downregulation of protein synthesis and an upregulation of protein degradation which is common in proteotoxic stress responses15. Chaperone proteins as a group tend to respond similarly to proteasome genes, with the exception of several chaperones that show a strong response in metagroup 3 (e.g. *DNAJA2* and *SPAG16*) suggesting that they might be involved in the folding of unfolded substrates in these compartments (Figure S14). A few proteostasis gene sets demonstrated tight compartment-specific regulation, such as the chaperone and folding enzyme subsets of ER proteostasis (Fig. 5D). When examining all genes involved in ER proteostasis, we again see strong upregulation in response to folding stress in the ER, but interestingly, to a lesser extent, also in response to mitochondrial proteotoxicity (Fig. 5E). Altogether, we show that our platform can produce rich datasets of proteotoxic stress across cellular compartments, unraveling shared and specific responses.

### Protein misfolding in the mitochondria directly induces the IRE1ɑ branch of the ER UPR

We closely inspected the crosstalk in stress responses seen between the ER lumen, mitochondrial matrix, and CYT_pqc. Mechanisms that could explain certain vectors of this crosstalk have been described in the past. For example, proteotoxic stress can lead to import of misfolded proteins into the mitochondria46, mitochondrial stress can be propagated to the cytosol47, and aggregation-prone proteins have been reported to translocate from the ER to the mitochondria as well48,49. Furthermore, proteotoxic stress from various sources, whether ER, mitochondrial, or cytosolic, can trigger a shared homeostasis pathway known as the integrated stress response (ISR)50,51. Proteotoxic signaling between the mitochondria and ER, particularly from the mitochondria to the ER, has not been well explored despite the extensive body of literature describing the interplay between the two organelles52. In our data, we noticed a marked induction of ER stress response genes, such as *CANX*, *UFD1*, and *DDOST*, upon protein misfolding in the mitochondria (Fig. 5E). Interestingly, certain canonical ER stress response genes, such as *HSPA5, HSP90B1*, and several protein disulfide isomerases (Fig. 5D, new supp), were exclusively upregulated in the ER tag group, suggesting a distinct form of the ER stress response to protein misfolding in mitochondria.

To validate the existence of an ER stress response signature in response to mitochondrial proteotoxic stress, we explored orthogonal published datasets in which mitochondrial proteostasis was perturbed. We analyzed data generated with a mitochondrial HSP90 inhibitor, GTPP, as well as a mitochondrial protease inhibitor, CDDO, to profile the mitochondrial unfolded protein response in mammalian cells53. In addition to mitochondrial stress genes, some of the most significant upregulated genes with brief treatment of either drug are part of the ER stress response (Fig. 5F). This phenomenon is also conserved in yeast as demonstrated by analysis of cells with a conditional knockout of the mitochondrial presequence protease MPP54. Two hour inactivation of MPP with a temperature-sensitive mutant subunit, *mas1ts*, resulted in consistent upregulation of ER stress-associated genes (Fig. S15A). Finally, to elucidate whether the mitochondria-induced ER stress response is activated at the ER as opposed to exclusively through a downstream signaling pathway such as the ISR, we introduced an XBP1 splicing reporter into HAP1 cells that detects the activity of the IRE1ɑ branch of the ER UPR20. We found in our cells that both GTPP and CDDO induce XBP1 splicing, and furthermore, GTPP-induced splicing exceeds that induced by ER calcium depletion with thapsigargin (Fig. 5G, Fig. S15B-C). These results indicate that proteotoxic stress in the mitochondria can directly perturb the ER, causing activation of ER stress sensors such as IRE1ɑ, inducing XBP1 splicing, and further upregulating various ER stress genes.

### Improved tagging and optical sequencing protocols to investigate poorly annotated genes in protein quality control

After exploring the relationships between tag group transcriptional responses, we next compared individual gene profiles with the aim of revealing coordinated modules associated with localized proteotoxic stress. We used minimum distortion embedding55 to visualize the similarities between transcriptional profiles of responsive genes and identify clusters that may represent potential proteotoxic response pathways (Fig. 6A). The position of the cluster on the embedding reflects specific response profiles: clusters towards the left hemisphere represent responses which are upregulated in metagroup 3 and downregulated in metagroup 1, and vice versa for the right hemisphere. The vertical axis of the plot, conversely, represents various compartment-specific patterns. Many clusters are enriched for known functional categories, such as translation and proteostasis. Notably, oxidative phosphorylation was anticorrelated with proteostasis responses, demonstrating upregulation in metagroup 3 and downregulation in metagroup 1. The full list of clusters, associated genes, and the top five functional enrichments per cluster can be found in Table 3. We next asked whether we could use the cluster association of poorly annotated response genes to infer their role in proteostasis. Because our data suggests that compartment-specific proteotoxic stress leads to upregulation of genes associated with that compartment (Fig. 4D, Fig. S11, Fig. S12), we hypothesized that many of the poorly annotated genes would exhibit localization patterns related to their response profiles.

**Figure 6.**
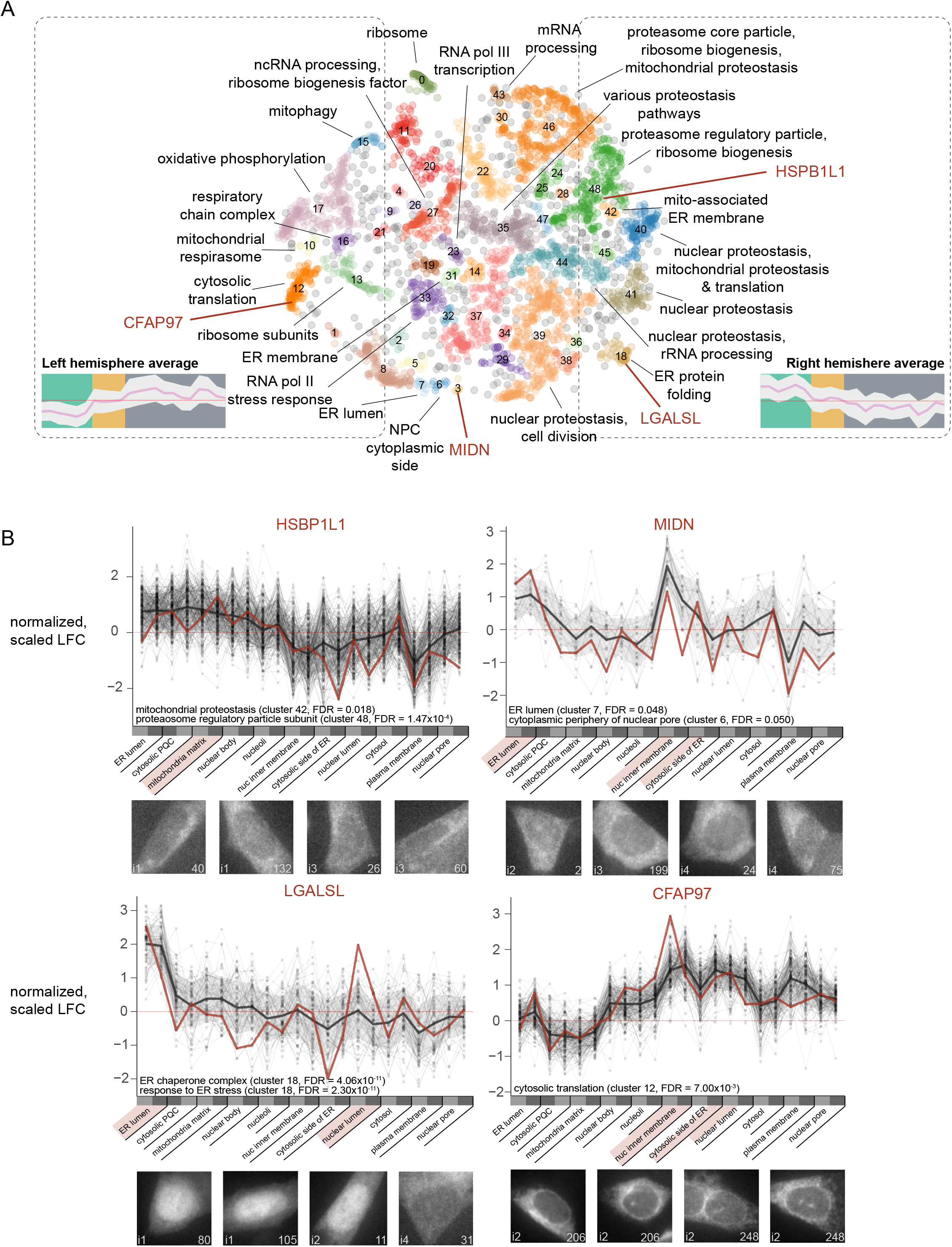
Unbiased clustering and functional annotation of proteotoxic stress-responsive genes. A) Minimum distortion embedding of gene responses to proteotoxic stress in different compartments. Each point is a gene colored by unbiased clustering, and enriched functional terms for each cluster are manually summarized in cluster labels. The mean and standard deviation Z-normalized log fold change across metagroups of left and right hemisphere genes is depicted. Select poorly-annotated genes are labeled in dark red. B) Poorly characterized proteins are implicated in protein quality control by their localization pattern and transcriptional response to protein misfolding across compartments. Proteins were endogenously tagged with second-generation SPOTLITES and their localization patterns identified by accelerated optical sequencing. Cell images showing localization patterns are annotated with the intron number on the bottom left of each image and the sgRNA ID on the bottom right. The expression pattern of the unannotated gene in response to hydrophobic tagging in each tag group is represented in dark red above the cell images, with all genes in the associated clusters in thin black lines and the average of those genes is represented by a thick black line. In the case of *MIDN*, small adjacent clusters 3, 6, and 7 were considered in aggregate, as *MIDN*’s cluster alone had too few genes for reliable recovery of functionally-enriched terms. Relevant functional enrichment terms associated with the cluster of each depicted gene are listed in the lower left-hand corner of the transcriptional profile plots.

To further investigate these genes and establish the utility of SPOTLITES as an efficient approach to explore endogenous function, we generated a SPOTLITES library using an improved protocol (Fig. 7A-B). To remove epigenetic barriers to tag insertion (Fig. 2F), we tested activating domains for their capacity to increase tagging efficiency (Fig. S17A). Fusing Cas9 to dMSK1, a hyperactive chromatin kinase mutant that phosphorylates histones56, improved the tagging success rate at all loci tested (Fig. S17B). Second, we capitalized on next-generation HaloTag fluorescent ligands with better signal to noise ratios and photostability57,58 for FACS-based purification and optical phenotyping (Fig. 7A, red text).

**Figure 7.**
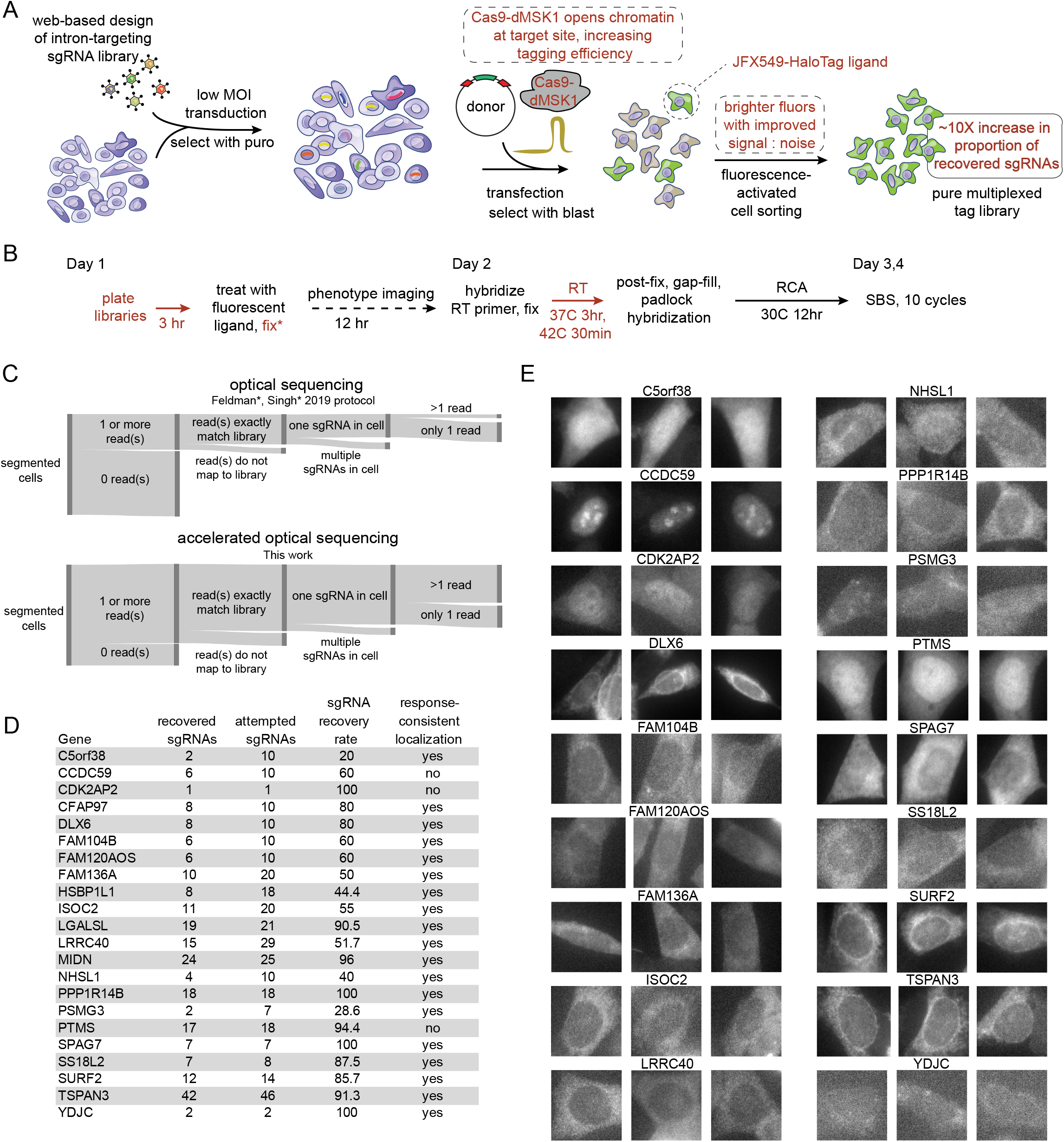
Improved tagging and optical sequencing protocols to investigate poorly annotated genes in protein quality control. A) Second-generation SPOTLITES tagging protocol and B) accelerated optical sequencing protocol with major changes highlighted in red text. C) A Sankey diagram quantitatively representing the proportion of cells segmented at 10X magnification that are lost at each step of filtering in the analysis of standard optical sequencing (top panel) and accelerated optical sequencing (bottom panel). D) Recovery rates of sgRNAs targeting poorly annotated proteins following second-generation SPOTLITES library generation and readout with accelerated optical sequencing. An sgRNA was considered ‘recovered’ if represented by at least 30 cells, each with two reads perfectly matching the expected sequence. The final column indicates whether tagged cells exhibited a localization pattern consistent with the compartments where hydrophobic tagging induces upregulation of the poorly annotated gene. E) Cell images of all poorly annotated proteins listed in subfigure D.

To improve cell recovery during optical sequencing36, we built on recently published advancements59 with three main modifications (Fig. 7B, red text). First, we began processing the library for optical phenotyping immediately after cells adhered in order to minimize the overlap between neighboring cells. Second, we incorporated 0.1% glutaraldehyde in the pre-reverse transcription fixation step to maximally immobilize gRNA-encoding mRNAs and potently inhibit endogenous RNases. Finally, we significantly abbreviated the reverse transcription reaction, shortening it from overnight to 3 hours at 37C followed by 30 minutes at 42C. These modifications to our tagging and optical sequencing pipeline, as well as additional improvements in library generation, contributed to a dramatic increase in the rate of cell recovery (Fig. 7C) and gRNA recovery (Fig. 7D), and significantly shortened the total bench time required for library generation and analysis to approximately six weeks.

With this improved protocol, we systematically compared the localization patterns of select poorly annotated response proteins to their gene clusters (Fig. 6A) and transcriptional stress response profiles (Fig. 6B). We found that proteins displayed localization patterns largely reflecting the compartments in which they were upregulated in response to protein misfolding (Fig. 6B; Fig. 7D-E). For example, *HSPB1L1* is predominantly upregulated in response to misfolding in the mitochondria and clusters with genes associated with regulation of the proteasome and near genes associated with mitochondrial proteostasis (Fig. 6A). Though not experimentally characterized, it exhibits a high degree of sequence homology to HSPB1, a small heat-shock protein that is upregulated to facilitate protein re-folding and degradation in the mitochondrion and other compartments60,61. Correspondingly, imaging endogenously tagged HSPB1L1 revealed a pattern consistent with a mitochondrial localization (Fig. 6B), which combined with the strong response to induced misfolding in the mitochondria, leads us to believe that HSPB1L1 may represent a novel mitochondrial heat shock protein.

Even in the ER, an organelle with well-characterized quality control systems, we found a number of poorly annotated genes that respond to protein misfolding. For example, *MIDN* is strongly upregulated in response to misfolding in the ER lumen and moderately upregulated in other compartments (Fig. 6B). Further, *MIDN* clusters near genes significantly enriched for ER-related GO:CC terms (Fig. 6B). Endogenously-tagged MIDN consistently localizes to the ER, with extra-ER cytosolic patterning and slightly brighter intensity along the nuclear membrane, thus mirroring its transcriptional signature. In a similar vein, *LGALSL* clusters with known genes involved in ER proteostasis (Fig. 6A). Closer investigation of the transcriptional responses reveals sensitivity to misfolding in both the ER and the nucleoplasm (Fig. 6B). Interestingly, LGALSL exhibits tag-site dependent localization: tags in two introns exhibit localization to both the ER and the nucleoplasm, while a third intron leads to exclusive ER localization (Fig. 6B, panel 3). While further investigations are required to determine the definitive steady state localization of the protein, our data show that it is capable of localization to either organelle and thus may have a direct role in ER and nuclear proteostasis. Two additional genes that were transcriptionally responsive to protein misfolding in the ER were overexpressed with C- and N-terminal tags and displayed localization to the secretory pathway, further supporting general concordance between transcriptional roles in stress and localization patterns (Fig. S16A).

Importantly, we can also investigate stress responsive proteins in compartments where protein quality control is very poorly understood. For example, *CFAP97* is acutely upregulated in response to protein misfolding at the nuclear membrane, and to a lesser extent, at the cytosolic face of the ER (Fig. 6B). CFAP97 is further implicated in protein quality control in these compartments by its inclusion in a cluster associated with cytosolic translation, which may point to a role in modulating translation at the nuclear membrane and ER in response to misfolding (Fig. 6A). Endogenously tagged CFAP97 localizes clearly to the nuclear membrane, and exhibits cytosolic patterning that may reflect localization to the cytosolic surface of the ER, consistent with its response to perturbation in those compartments.

Our data demonstrate the potential of SPOTLITES libraries to explore localization patterns, measure responses to direct perturbation of tagged proteins, and infer novel functions of poorly annotated proteins at scale. Altogether, by combining web-based construction of an intron tagging library (www.pooledtagging.org) with improved and expedited protocols, we provide an efficient and powerful end-to-end approach for imaging and perturbation of proteome dynamics.

## Discussion

We developed SPOTLITES (Scalable POoled Targeting with a LIgandable Tag at Endogenous Sites) to address the lack of high throughput approaches available to study the proteome using both imaging and direct protein perturbation. Our approach enables the efficient generation of complex pooled cell libraries with a wide variety of endogenously tagged proteins using multiple tag sites to address the possible interference of the tag on the targeted endogenous protein. Using HaloTag as our handle, we can manipulate and visualize the proteome using any of a suite of cell-permeable HaloTag ligands8,13,62 or ligand-orthogonal tools such as HaloTag antibodies for combining both imaging and perturbations. We used optical sequencing followed by image analysis to generate a detailed localization map for multitudes of proteins in a single experiment. The multi-functionality of HaloTag enabled us to use the same cell pool for unbiased mapping of protein misfolding responses.

Proteotoxic stress can be achieved by labeling HaloTag, particularly the metastable HaloTag2, with hydrophobic tags that chemically destabilize the protein23. This approach allows us to measure how the local proteostasis machinery responds to a sudden onset of misfolded protein, avoiding the off-target effects and toxicity associated with common pharmacological methods, for example by inhibiting N-linked glycosylation or calcium uptake in the ER20. Our findings reveal that compartments exhibit distinct transcriptional responses to proteotoxic stress (Fig. S12). In the ER, differentially expressed genes are very well characterized, while in other compartments, they may represent novel proteostasis responses. There is also notable similarity between certain compartment responses, particularly in the differential expression of proteostasis-related genes which may indicate a general response or direct crosstalk (Fig. S13; Fig. 5B-E). Given the interconnected nature of the cell, considerable crosstalk is not surprising, and in fact various aspects of organelle crosstalk have been described46–48,63. We highlight the ability of protein misfolding in the mitochondria to induce ER stress (Fig. 5E-G). Both the ER and the mitochondria have mechanisms to initiate the aptonymous integrated stress response which may account for some of this crosstalk50,51. However, we find that proteotoxic stress in the mitochondria can induce XBP1 splicing through the ER-localized stress sensor IRE1α (Fig. 5G), which may be mediated by mitochondria-associated ER membranes (MAMs) at which IRE1α has been observed to accumulate64,65. Other cellular compartments, such as the nucleoplasm and specific regions of the cytosol, do not seem to elicit a canonical proteostasis response upon protein destabilization. In these compartments, proteostasis appears to be downregulated while translation and oxidative phosphorylation are upregulated (Fig. 5C; Fig. 6A). The exact implications of these activities remain unclear, but they have been associated with cell death potentially through oxidative stress66, indicating a possible strategy to impede cell growth when proteostasis is not available.

While we discovered many new response genes to localized protein misfolding, our curated dataset contains fewer than 40 tagged proteins spanning many localization patterns. The limited scale was primarily driven by the skew of sgRNAs within the pooled library, resulting in a very large number of cells required for covering low abundance sgRNAs with sufficient coverage. The skew is primarily a result of differential tagging efficiency across genomic locations, which our research, as well as that of others, suggests is associated with different epigenetic states of the targeted sites (Fig. 2)33,34. To address this issue, our second-generation SPOTLITES protocol uses Cas9 fused to a chromatin modifying domain, dMSK1, that consistently increases tagging efficiency at multiple sites (Fig. S17). Grouping validated sgRNAs based on empirical or predicted tagging efficiency is another strategy to minimize skew in second generation libraries. Future libraries will also capitalize on empirically-validated tag sites to dramatically reduce the number of sgRNAs required for each protein. This increased scale will enable the generation of more detailed atlases of localization patterns and transcriptional responses to direct protein perturbation.

One clear limitation of any tagging approach, whether endogenous or ectopic, is the possibility of disrupting localization and function as a consequence of protein fusion. For example, tagging can affect localization signals, regulatory regions, and interaction sites. Previous attempts at large-scale C- and N-terminal arrayed tagging libraries were unable to tag ∼25% of attempted genes10. Without any prior knowledge, internal tagging may be more likely to disrupt protein structure, yet the scalability of our approach enables us to test a large number of tagging sites and recover functional tags for the majority of genes. However, because our method relies on introns, which are absent from 16% of human genes, it is not universally applicable to all protein coding genes. Thus a combination of tagging approaches, both internal and terminal, is necessary to achieve comprehensive and concise characterization of the proteome. Development of smaller fusion tags, together with advanced computational structure prediction methods67,68, will further enable the generation of large, minimally disrupted libraries.

The scalability of the pooled approach enables the generation of detailed cellular organization maps that could be measured across many environmental conditions. Changes in individual protein subcellular localization and abundance in response to stimuli represent a unique high-dimensional single cell readout for the discovery of molecular mechanisms not reflected in protein or mRNA levels alone. With large enough libraries, even subtle subcompartment domains can be reliably distinguished and their proteomes annotated. In addition to fluorescent and hydrophobic ligands, HaloTag can be labeled with E3 ligase-recruiting degraders69 and general inducers of protein-protein interactions70,71, that when applied on SPOTLITES libraries expand functional genomics into the protein space with a variety of direct perturbation modalities. Altogether, we provide an end-to-end platform for the generation and analysis of pooled cell libraries with endogenously tagged proteins, accompanied by a website for easy library design (www.pooledtagging.org), to facilitate powerful and highly parallelized investigations of protein regulation and dynamics across environments, perturbations, and cell types.

## Acknowledgements

SES and YVS have equally contributed to this work and their names are listed in alphabetical order. We would like to thank the Shalem lab for extensive discussions related to this manuscript. We would also like to thank members of the CHOP High Throughput Sequencing Core and Single Cell Technology Core, including Dr. David Smith for helpful discussions on data processing. This work was supported by the following grants: DP2GM137416 from NIH/NIGMS, SAP#4100083086 from PA DoH and R03NS111447-01 from NINDS awarded to OS. F32CA239499 from NCI and K99AG075256 from NIA awarded to YVS. T32GM008216 from NIGMS and F31HG011185 from NHGRI awarded to SES. R35GM142505 from NIGMS awarded to GMB.

## Author contributions

SES, YVS, and OS conceived the study, designed the experiments, and interpreted experimental data. YVS and SES collaborated to perform all experiments in the manuscript except those noted below. YVS performed biochemistry assays, optimized tagging efficiency, and generated and analyzed the bulk sequencing and single-cell RNA sequencing data. SES performed, optimized, and analyzed the high throughput microscopy and optical sequencing experiments. SES and TL wrote code for image processing and cross-objective mapping. TL performed the deep learning image analysis, including writing the convolutional neural network classifier, and training and applying the cytoself algorithm on HAP1 cell images. TL generated and analyzed the HeLa tagged library. GMB synthesized the HyT36 ligand and provided quality control. OS supervised the work. SES, YVS, and OS wrote the manuscript.

## STAR METHODS

## KEY RESOURCES TABLE

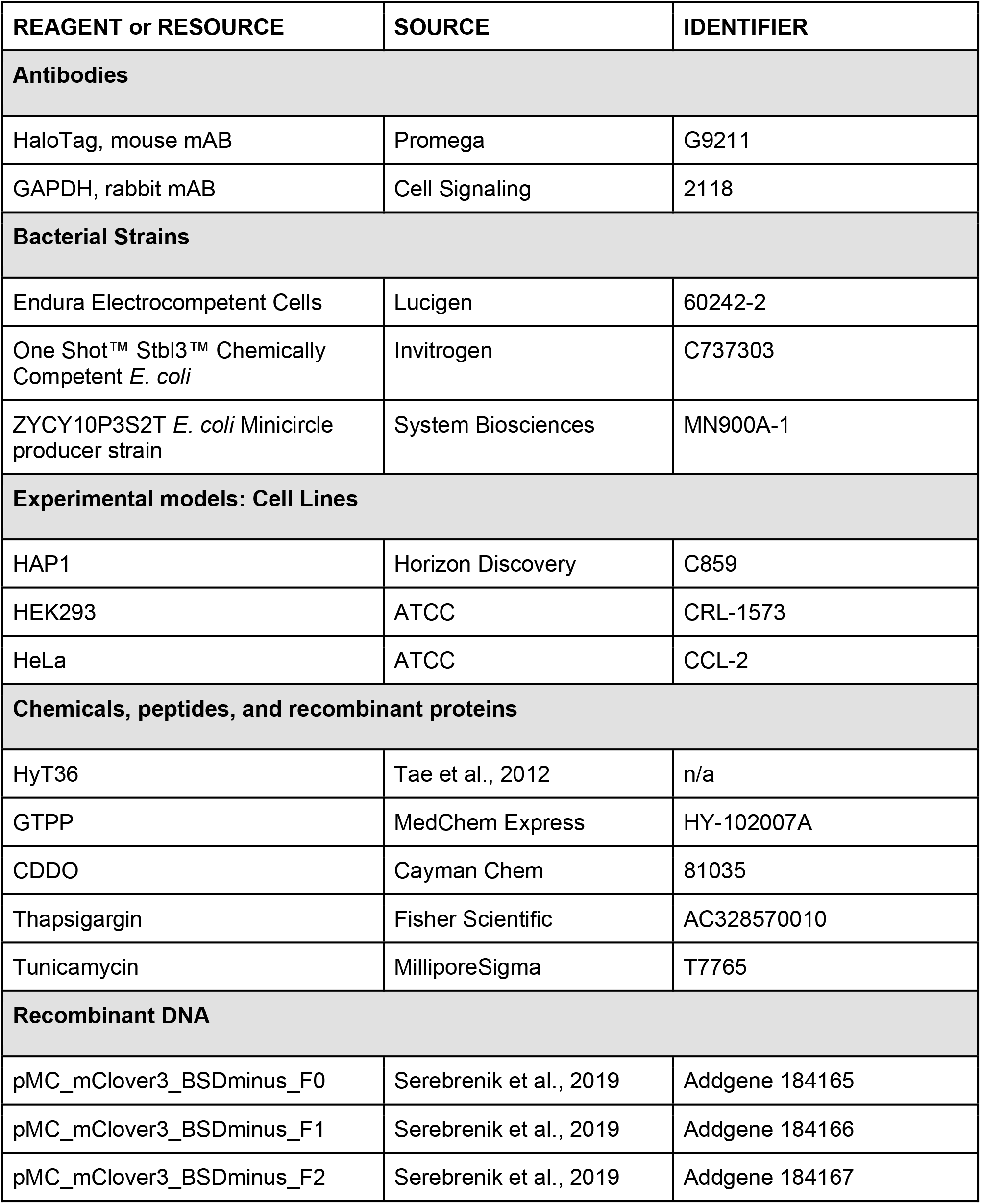
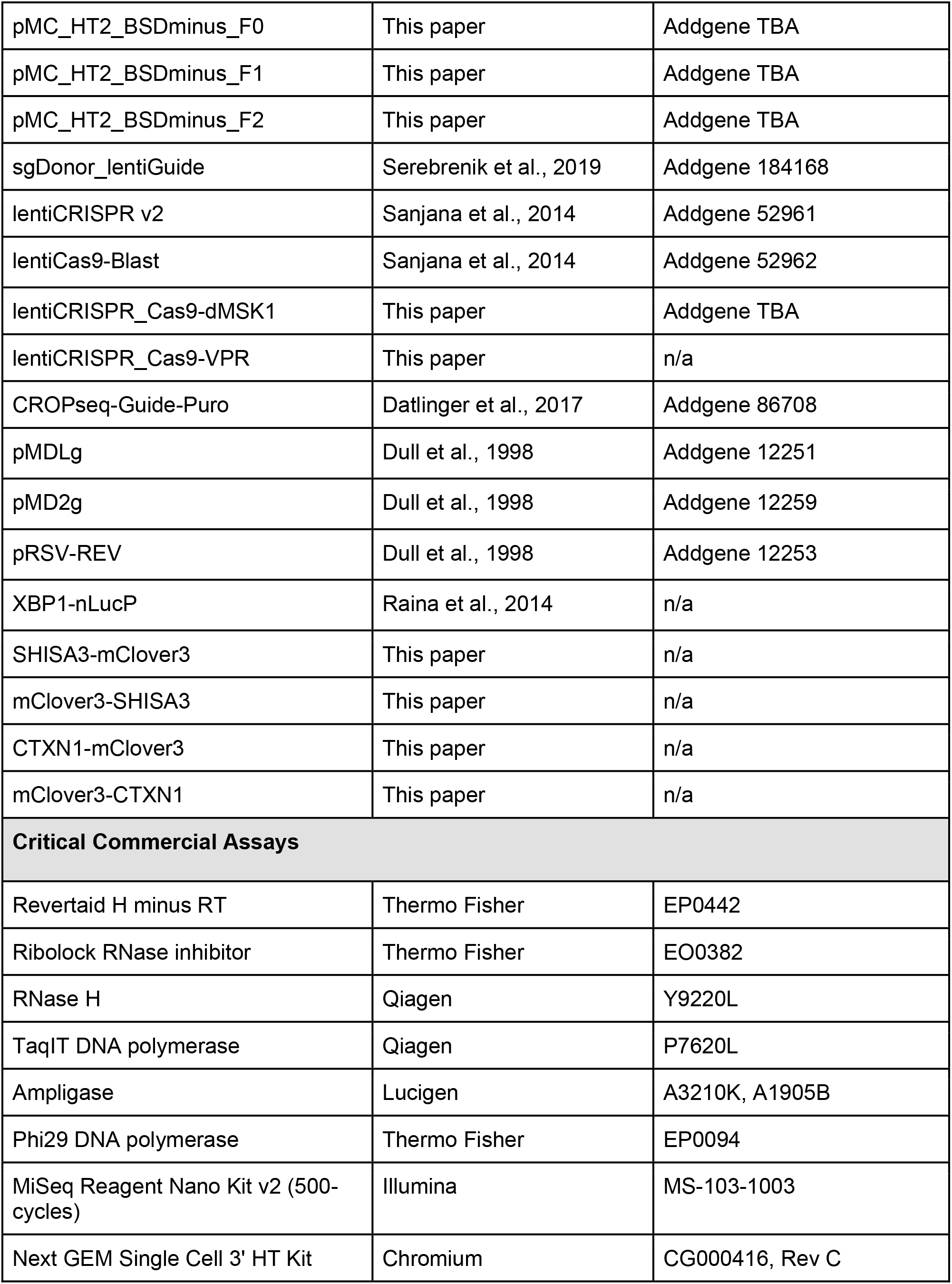
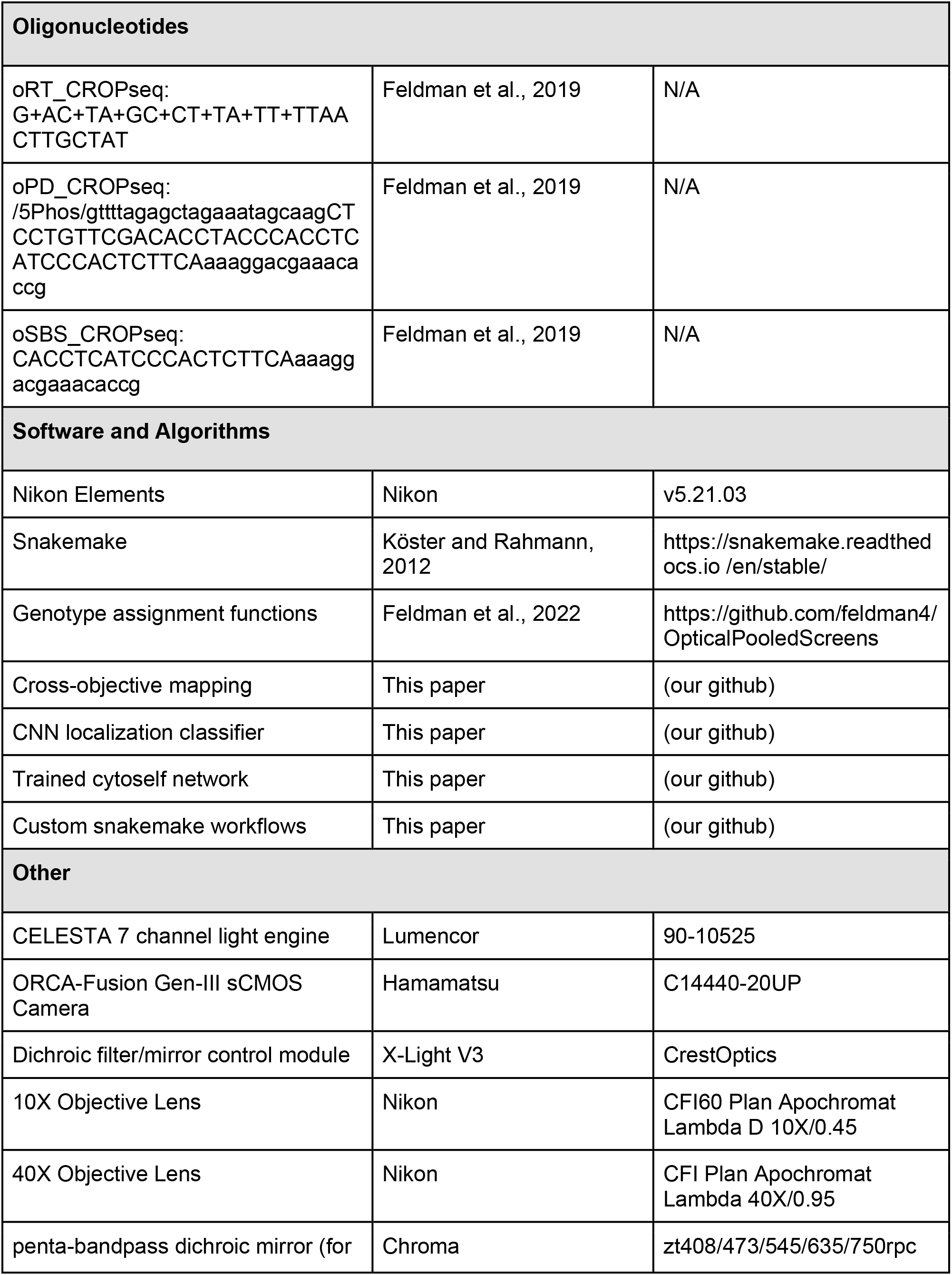
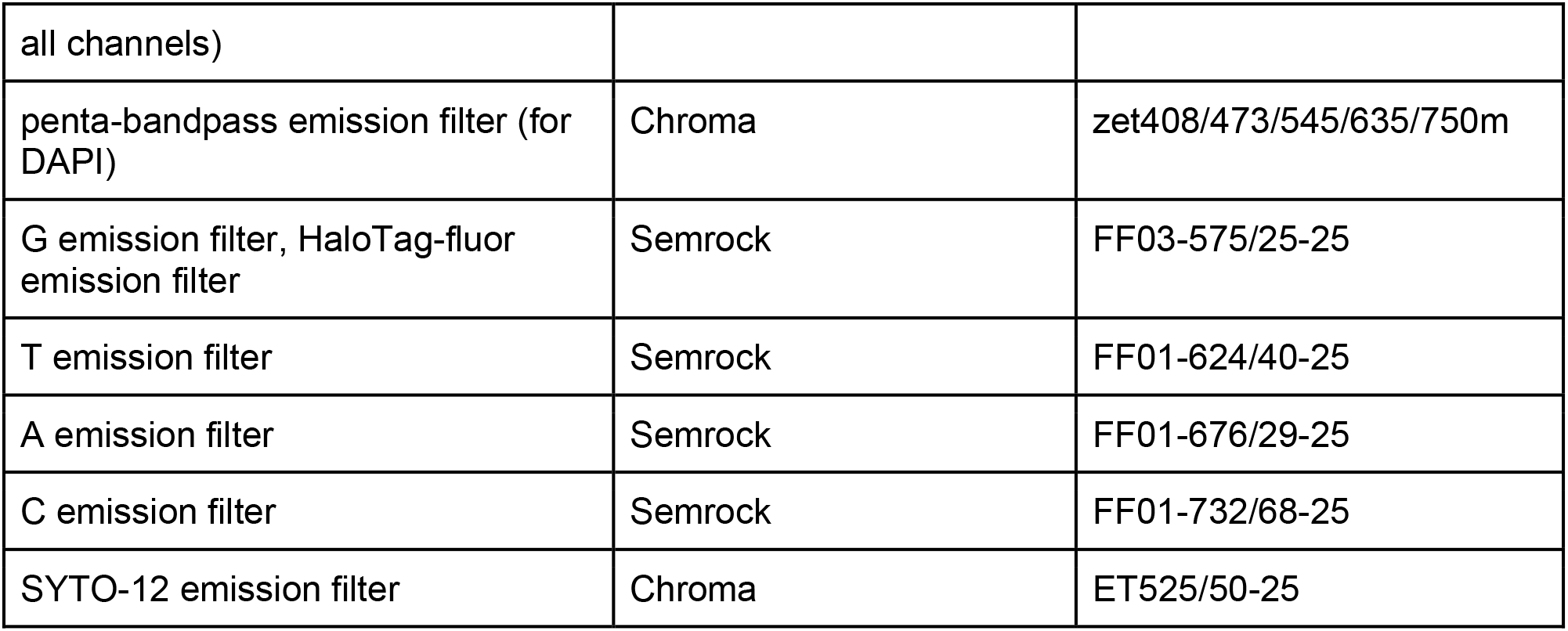

## LEAD CONTACT AND MATERIALS AVAILABILITY

### Lead contact

Further information and requests for resources and reagents should be directed to and will be fulfilled by the lead contact, Ophir Shalem.

### Materials availability

Materials used in this study will be provided upon request and available upon publication, all plasmids will be deposited on AddGene and library design can be performed freely on www.pooledtagging.org.

### Data and code availability

- Raw sequencing data will be deposited on GEO to be available upon paper publication.
- All code for the analyses performed will be publicly available on github.
- Cell albums for all tagged proteins will be available on a public web site.

## EXPERIMENTAL MODEL AND SUBJECT DETAILS

### Tissue Culture

Cell lines, including HAP1 (Horizon Discovery), HEK293 (ATCC CRL-1573), and HeLa cells (ATCC CCL-2), were handled according to manufacturer’s instructions. Clonal HAP1 haploid lines were generated by single cell sorting and genome editing was performed within 1-2 weeks of expansion to ensure most cells were haploid during editing. HAP1 cells were cultured in IMDM (Gibco, 12440053) + 10% fetal bovine serum (Thermo Fisher Scientific, FB12999102) + 1% antibiotic-antimycotic (Gibco, 15240062). HEK293 and HeLa cells were cultured in DMEM (Gibco, 11995065) + 10% fetal bovine serum (Thermo Fisher Scientific, FB12999102) + 1% antibiotic-antimycotic (Gibco, 15240062). All cells were dissociated with TrypLE Express (Thermo Fisher Scientific, 12605010).

## METHOD DETAILS

*Pooled tagging: plasmid library generation*

In generating pooled tag cell libraries, the protocol for “plasmid library generation” and “viral library generation and transduction” (next section) follows typical CRISPR library generation methods72. Nonetheless, we present all steps in detail below.

1. An oligo pool was synthesized by Twist Bioscience or as oPools Oligo Pools (IDT) with the following design logic:

5’ - fw. primer (N20) - BsmBI binding (CGTCTC) - backbone overhang (ACACCG) - spacer seq. (N20) - backbone overhang (GTTTT) - BsmBI binding (GAGACG) - rev. primer (N20) - 3’

2. Oligos were PCR amplified and purified. 25 μl reactions were prepared for each sublibrary with the KAPA HiFi PCR Kit (Roche, KK2502) containing 0.5 μl of the oligo pool resuspended to 20 ng/μl, 5 μl 5x KAPA HF buffer, 0.75 μl 10 mM dNTP mix, 0.5 μl polymerase, 0.75 μl appropriate 10 μM primer mix. PCR cycling parameters: 1) 3 min. at 95°C, 2) 20 sec. at 98°C, 3) 15 sec. at 65°C, 4) 15 sec. at 72°C, 5) repeat steps 2-4, 10x, 6) 1 min. at 72°C. PCR products were gel purified with the Monarch Gel Extraction Kit (NEB, T1020).

3. Amplified oligos were ligated into an sgRNA-expressing backbone by Golden Gate cloning with BsmBI (NEB, R0739) and T7 ligase (NEB, M0318), using 5 ng of PCR product and 50 ng of the CROPseq backbone (Addgene, 86708). PCR cycling parameters: 1) 5 min. at 42°C, 2) 5 min. at 16°C, 3) repeat steps 1-2, 50x, 4) 10 min. at 65°C.

4. Plasmids are transformed and purified. Golden Gate products were first concentrated by IPA precipitation. 1 volume sample was mixed with 1 volume isopropanol in 50 mM NaCl and incubated at room temperature for 15 minutes. Samples were then centrifuged at 15 k x g for 15 minutes at room temperature and washed 2x with cold 70% EtOH. After drying the DNA pellets, samples were resuspended in 5 μl H_2_O. Electroporation was performed according to manufacturer’s instructions into Endura ElectroCompetent Cells (Lucigen, 60242-2). Dilution of transformed cells and estimation of the number of colony forming units allowed for calculation of plasmid library coverage, which ranged from 500x to 10k x depending on the library size. Bacteria was cultured overnight and plasmids were purified with a maxiprep kit according to manufacturer’s instructions (Thermo Fisher Scientific, A31217).

### Pooled tagging: viral library generation and transduction

1. Lentivirus was produced from HEK293T cells seeded into a 15 cm tissue culture dish at 10 M cells / dish. Transfection was performed with PEI (Polysciences, 24765) in Opti-MEM (Thermo Fisher Scientific, 31985062). 136.35 μl PEI (3 μl PEI : 1 μg DNA) was incubated in 1.25 ml Opti-MEM, and a separate solution of 1.25 ml Opti-MEM was mixed containing: 1) 20 μg plasmid pool, 2) 13.25 μg pMDLg, 3) 7.2 μg pMD2g, 4) 5 μg pRSV-REV. Both solutions were combined, incubated for 15 minutes, and 2.5 ml added dropwise to cells containing 15 ml of antibiotic-free media. Media was replenished the following day, and viral aliquots were collected 48 hours post-transfection, filtered through 0.45 μm cellulose acetate filters (VWR, 76479-040) or 0.45 μm PES filters (Thermo Fisher Scientific, 50-607-518), and frozen at -80°C.

2. Virus was titered and cells were transduced. Titering was performed by puromycin selection and cell counting. Our initial pooled tagging library (Fig. 1) was generated by infection with the lentiviral library at ∼30% transduction efficiency and in amounts to achieve 1000 transduced cells per sgRNA. To further minimize the proportion of cells with more than a single transduction, subsequent libraries were infected at a transduction efficiency < 10% and in amounts to achieve 5 k transduced cells per sgRNA. Virus was administered by “spinfection”, where cells and virus were mixed with polybrene (Millipore TR-1003-G) in antibiotic-free media at 2 M cells per well of a 12-well plate. Plates were spun at 32°C for 1 hour at 1000 x g and placed in a tissue culture incubator overnight. The following day, cells were expanded into 15 cm dishes in the presence of 1 μg/ml puromycin (Gibco, A1113803), and selection took place over 5 days.

### Pooled tagging: tagged cell library generation

1. Transduced cells were transfected with reagents necessary for homology-independent tagging. Based on an assumed average tagging efficiency of 0.05% in HAP1 cells, we transfected enough cells for 10 - 200x coverage of the tagging library, depending on the library size. Cells were plated at ∼13 M cells per 15 cm dish in 15 ml of antibiotic-free media. Transfection was performed with Lipofectamine 3000 (Invitrogen, L3000001). 89 μl Lipofectamine 3000 reagent (3 μl reagent : 1 μg DNA) was incubated in 1.25 ml Opti-MEM, and a separate solution of 1.25 ml Opti-MEM was mixed containing: 1) 8.6 μg tag donor pMC_*X*_BSDminus_*Y* (where *X* refers to the fusion domain and *Y* refers to the sequence phase of the library), 2) 7.6 μg sgDonor_lentiGuide, 3) 13.5 μg lentiCRISPR, 4) 60 μl P3000 reagent (2 μl reagent : 1 μg DNA). The molar ratio of tag donor to any other plasmid was kept 5 : 1. The tag donor was also introduced as a minicircle generated with the MC-Easy Minicircle DNA Production Kit according to manufacturer’s instructions (System Biosciences, MN920A-1). Both solutions were combined, incubated for 15 minutes, and added to cells dropwise. Media was replenished after 5 - 6 hours of incubation.

2. Cells with successful genomic integration of the tag donor were selected using the blasticidin resistance cassette11. 48 hours after transfection, media with 15 μg/ml blasticidin (Gibco A1113903) was added to cells. Cells were incubated for 3 days until all untransfected cells depleted. Cells were subsequently expanded for 3 days in the presence of 5 μg/ml blasticidin. After expansion, cells were split in the presence of 15 μg/ml blasticidin and another round of selection occurred, this time between cells with and without genomic integration of the donor. Again, cells were expanded in the presence of 5 μg/ml blasticidin until they were ready for sorting.

3. Cells with expressed endogenous fusion proteins were purified by fluorescence-activated cell sorting (FACS) as described in the “Flow cytometry” section. Prior to sorting, HaloTag was labeled with HaloTag-TMR (Promega, G8251) as described in the “Cell culture” section. 1 - 5 M cells per library were sorted. This amounted to 15 - 25 k x coverage for our smaller libraries (Figs. 4 & 7), and at least 2000x average coverage for our initial tagging library based on the sgRNA success rate (Fig. 1). A second round of FACS was performed for each library to ensure the percentage of HaloTag-expressing cells was at least 95%.

### Pooled tagging: tagged library amplicon sequencing

Tag library composition was determined by either *in situ* sequencing (STAR Methods) or bulk sequencing of amplicons. For bulk sequencing, genomic DNA was extracted by incubation with 6 ml of lysis buffer (50 mM Tris pH 8.0, 50 mM EDTA, 1% SDS, 0.1 mg/ml proteinase K (Qiagen, 19131)) at 55°C for 16 hours. RNA was digested with 0.05 mg/ml RNAse A (Qiagen, 19101) added to the buffer and incubated at 37°C for 30 minutes. Protein was precipitated after chilling samples on ice and adding cold ammonium acetate (Sigma, A1542) to a final concentration of 1.875 mM. Samples were vortexed for 20 seconds and centrifuged for 20 minutes at 4,300 x g. The supernatant was isolated and gDNA was precipitated with isopropanol. Pellets were resuspended in 500 μl nuclease-free water at 65°C for 1 hour and at room temperature for 16 hours. Extracted gDNA was PCR amplified (Takara, RR001A) using a 30-cycle protocol. Each reaction had 5-10 μg gDNA template. PCR products were purified by gel extraction (NEB, T1020) and sequenced on a NextSeq 500 (Illumina) with a 5% spike-in of PhiX v3 (Illumina FC-110-3001).

### CRISPR KO screen: library generation, lentivirus production, screening, genomic DNA isolation, and sequencing

CRISPR KO screening for growth-essential genes was performed as previously described6,73 with minor modifications. Briefly, a targeted oligo pool derived from the Brunello CRISPR-KO library73 was synthesized by Twist Bioscience. The oligo pool was PCR amplified according to manufacturer’s instructions (Roche, KK2502), purified by gel extraction (NEB, T1020), and cloned into the CROPseq-Puro (Addgene) backbone by Golden Gate cloning with BsmBI (NEB, R0739) and T7 ligase (NEB, M0318). The product was precipitated with isopropanol and resuspended in 3 μl nuclease-free water (Invitrogen, 10977015), followed by electroporation into Endura ElectroCompetent Cells (Lucigen, 60242-2) at ∼100x coverage. Plasmids were purified using a maxiprep kit (Thermo Fisher Scientific, A31217).

Lentivirus was produced from HEK293T cells seeded into a 15 cm tissue culture dish at 10 M cells / dish. Transfection was performed with PEI (Polysciences, 24765) in Opti-MEM (Thermo Fisher Scientific, 31985062). 136.35 μl PEI (3 μl PEI : 1 μg DNA) was incubated in 1.25 ml Opti-MEM, and a separate solution of 1.25 ml Opti-MEM was mixed containing: 1) 20 μg plasmid pool, 2) 13.25 μg pMDLg, 3) 7.2 μg pMD2g, 4) 5 μg pRSV-REV. Both solutions were combined, incubated for 15 minutes, and 2.5 ml added dropwise to cells containing 15 ml of antibiotic-free media. Media was replenished the following day, and viral aliquots were collected 48 hours post-transfection, filtered through 0.45 μm filters (VWR, 28145481), and frozen at -80°C.

Cas9-expressing HAP1 cells were generated by low multiplicity of infection transduction with lentivirus generated from lentiCas9-Blast74 (Addgene 52962) as described in the preceding paragraph, followed by 5 days of 10 μg/ml blasticidin selection and sorting-based purification of haploid cells (‘*DNA content analysis’*). Haploid Cas9-expressing HAP1 cells were infected with the lentiviral library at ∼35% transduction efficiency and in amounts to achieve 1000 transduced cells per sgRNA. Virus was administered by “spinfection”, where cells and virus were mixed with polybrene (MilliporeSigma, TR1003G) in antibiotic-free media at 2 M cells per well of a 12-well plate that had been coated with 0.1% gelatin (Sigma, G2500) for 15 minutes. Plates were spun at 32°C for 1 hour at 1000 x g and placed in a tissue culture incubator overnight. The following day, cells were expanded into 15 cm dishes in the presence of 1 μg/ml puromycin, and selection took place over 3 days. After puromycin selection, cells continued to be passaged every ∼3 days for two weeks in the presence of 10 μg/ml blasticidin to maintain Cas9 expression, with samples collected for potential processing at each passage. Genomic DNA was extracted and prepped for bulk sequencing as described in the section: *Pooled tagging: tagged library amplicon sequencing*.

### Bulk RNA sequencing

Cell pellets were frozen on dry ice and submitted for standard RNA sequencing by Azenta Life Sciences. RNA was purified by PolyA selection and libraries were prepared using the NEBNext Ultra II RNA Library Prep Kit (NEB, E7770). Samples were sequenced on a NextSeq 2000 (Illumina).

### HaloTag ligand synthesis and treatment

Subconfluent HaloTag-expressing cells were labeled with HaloTag-TMR (Promega, G8251), JFX-554 (Promega, CS315101), or JFX-549 (Janelia materials) at 20 nM final concentration for 15 minutes at 37°C with 5% CO_2_. Cells were then washed 3× with fresh media and incubated for an additional 30-60 minutes in media. Labeled cells were analyzed either by microscopy or flow cytometry.

HyT36 was synthesized as described and solubilized in DMSO75. For treatment, cells were replenished with media containing indicated concentration of HyT36 and incubated for the indicated lengths of time.

### Pharmacologic stress induction

ER and mitochondrial stress experiments were performed with 0.5 µM thapsigargin, 10 µg/ml tunicamycin, 10 µM GTPP, or 2.5 µM CDDO for indicated lengths of time. All were dissolved in DMSO which was at 0.1% final concentration.

### Flow cytometry and cell sorting

Cultured cells were trypsinized, resuspended in DMEM to ∼1×106 cells/ml, and filtered through a cell strainer. Cellular fluorescence was measured on a BD FACSAria Fusion (BD Biosciences). All cells were analyzed or sorted using an 85 µm nozzle. HaloTag-TMR fluorescence was detected by the 561 nm laser and the 582/15 filter. GFP fluorescence was detected by the 488 nm laser and filters 502LP and 530/30. Hoechst signal, or otherwise “autofluorescence”, was detected by the 405 nm laser and the 450/50 filter. Data were analyzed using the R package *flowCore* (v2.2.0). Polyclonal sorting was performed according to cytometer instructions into 15 ml conical tubes at 4°C partially filled with media. To culture sorted cells, tubes were centrifuged at 300 g for 3 min and the cells were resuspended in fresh media and then transferred to a tissue culture plate. Single cell sorting was performed directly into 96-well tissue culture plates containing 50 µl media.

### Confocal Microscopy

Cells were grown on coverslips and directly fixed in 4% formaldehyde (Electron Microscopy Sciences) in PBS (Thermo Fisher Scientific). Fixed cells were washed in PBS and coverslips were mounted on microscopy slides in ProLong Glass Antifade Mountant with NucBlue Stain mounting medium (Thermo Fisher Scientific, P36985). Images were acquired on a Leica TCS SP8 confocal microscope. Z-stacks (0.6 μm slices) spanning the entire volume of the cells were recorded with oil-immersion 63× Plan-Apochromat lenses, 1.4 NA.

### DNA content analysis

Cells were cultured in media containing 5 µg/ml Hoechst 33342 Solution (Thermo Fisher Scientific, 62249) at 37°C for 30-60 minutes. Cells were then immediately lifted and resuspended in PBS on ice. DNA content was measured using the flow cytometer with the 405 nm laser and the 450/50 filter.

### Immunoblotting

Cultured cells were pelleted, washed with PBS, and resuspended in RIPA lysis buffer (Cell Signaling, 9806) with 1× protease inhibitor cocktail (MilliporeSigma, P8340). Samples were normalized by bicinchoninic acid (BCA) assay (Cell Signaling, 7780), and loaded on a precast SDS-PAGE gel (Bio-Rad, 4561086). Immunoblotting followed using standard protocols. Imaging of blots was performed on a LI-COR Odyssey instrument. The following antibodies were used: HaloTag mAB (Promega, G9211, 1:500), GAPDH (Cell Signaling 2118, 1:2000), IRDye 680LT Goat anti-Rabbit (LI-COR 926-68021, 1:10,000), IRDye 800CW Goat anti-Mouse (LI-COR 926-32210, 1:10,000).

### Viability assays

CellTiter-Glo (Promega, G9241) was performed in black opaque 96-well tissue culture plates (Corning, 3596). The cell plate and reagent were equilibrated to room temperature for 30 minutes. 1 part reagent was added to two parts cell volume and the plate was mixed on an orbital shaker for 2 minutes followed by 30 minutes at room temperature. Luminescence was measured on a SpectraMax i3 plate reader.

The MTS or CellTiter 96 AQueous One Solution Cell Proliferation Assay (Promega, G3580) was performed in transparent 96-well tissue culture plates (Corning, 3596). The reagent was warmed to room temperature and 20 µl was added to cells in 100 µl media, which were then incubated in a tissue culture incubator for 1 to 4 hours. Absorbance at 490 nm was then measured on a SpectraMax i3 plate reader.

### Luciferase assay

Cells expressing an XBP1-nLucP reporter were cultured in black opaque 96-well tissue culture plates (Corning, 3596). Luciferase levels were detected using the Nano-Glo Luciferase Assay System (Promega, N1110). The cell plate and prepared reagent (1 part reagent : 50 parts buffer) were equilibrated to room temperature for 5-10 minutes. 30 µl of prepared reagent was added to 70 µl cell volume and mixed on an orbital shaker. Luminescence was measured after 3-5 minute incubation at room temperature on a SpectraMax i3 plate reader.

### High throughput phenotyping and in situ sequencing

For standard optical sequencing runs (Fig. 3, Fig. 5), HAP1 library cells were plated on glass-bottom 6 well plates (CellVis, P06-1.5H-N) 16 hours prior to treatment at 5×105 cells per well. Cells were deposited by adding 2 mL of cell suspension dropwise from a serological pipette moving in circles above the well. The next day, cells were incubated in 1.5 mL per well of 20 nM TMR HaloTag ligand (Promega, G8251) in pre-warmed medium for 15 minutes, washed 1X in 2 mL pre-warmed medium per well, and washed a third time for 45-60 minutes in the incubator to efflux unbound ligand. The third wash doubled as a treatment time for segmentation stain SYTO-12 (Invitrogen, S7574) at 100 nM. All washes were performed gently to avoid sloughing HAP1 cells, which tend to be rounded and not particularly well-adhered to the glass surface.

After the third wash/SYTO-12 treatment, media was aspirated at the bench and cells were processed as previously published25,36. In brief, cells were fixed in 4% formaldehyde (Electron Microscopy Services, 15710) in RNase-free PBS (ThermoFisher, AM9625) for 20 minutes, washed with RNase-free PBS 3X, and incubated for 30 minutes in 200 ng/mL DAPI (ThermoFisher, 62248) and 0.8 U/uL RiboLock (ThermoFisher, EO0382) in 2 mL RNase-free 2X SSC (ThermoFisher, AM9763) per well prior to phenotyping imaging.

For accelerated optical sequencing runs (Fig. 7), cells were plated on fibronectin-coated glass-bottom 12-well plates (CellVis, P12-1.5H-N). 150 uL of fibronectin (Corning, 354008) resuspended to 175 ug/mL in sterile water was ejected onto each well, and the plate was tapped to distribute the beaded liquid to the periphery of the wells. The plate was tilted and all liquid removed. The plate was cured at room temperature in the tissue culture hood for 45 minutes while cells were lifted. Cells were plated dropwise in a relatively large volume of medium (3 mL per well of 12w plate, 2.5×105 per well) and allowed to adhere for 45 minutes undisturbed before the plate was transferred to the incubator, as recently reported59. Cells were monitored under a brightfield microscope at 30-minute intervals and treated with HaloTag-fluor once approximately 80% of cells had adhered, which occurred three hours after plating. Cells were stained as in previous optical sequencing runs per ‘*HaloTag ligand synthesis and treatment’*, with one exception: JFX549-HaloTag ligand (Janelia materials) at 20 nM in DMSO was substituted for TMR HaloTag ligand. Following treatment with JFX549-HaloTag ligand and SYTO-12, medium was aspirated and cells were fixed as reported above, with the addition of 0.1% glutaraldehyde (Electron Microscopy Services, 16120).

Fixed cells in 2X SSC + RiboLock at 1:50 (0.8 U/uL) were imaged on an inverted Ti2 Nikon Eclipse microscope outfitted with an X-Light V3 (CrestOptics) for control of filters and dichroic mirrors (Key Resource Table) and a CELESTA 7 channel light engine (Lumencor, 90-10525) for sample illumination. Phenotyping images were collected on an ORCA-Fusion Gen-III sCMOS camera (Hamamatsu, C14440-20UP) with channel-specific filters (Key Resources Table) at 40X magnification (Key Resources Table) using the following run-specific settings: DAPI: 408 nm laser, 10% power, 10 ms exposure time; SYTO-12: 477 nm laser, 15% power, 200 ms exposure time; and HaloTag-fluor: 546 nm laser, 20% power, 200 ms exposure time. Exhaustive tiling of the well was achieved via an automated JOB programmed in Nikon Elements (v5.21.03).

Imaging four wells of a 12-well plate with these settings required approximately 12 hours. After imaging, standard optical sequencing samples were processed according to previously published protocols25. For accelerated optical sequencing, recent advancements59 were incorporated along with additional modifications that have not been previously reported to our knowledge. In brief, each well was washed 2X with 1 mL RNase-free PBS + 0.05% Tw (‘PBST’), then dehydrated in 0.5 mL prepared 70% ethanol at room temp for 30 minutes. Ethanol was replaced with PBST in three successive washes where 2.5 mL PBST was added and all but 0.5 mL of liquid was removed, followed by a fourth wash where all liquid was replaced with PBST. 250 uL per well of 1 uM reverse transcription primer in PBST was hybridized for 30 minutes at room temperature, followed by fixation in freshly-prepared 3% formaldehyde and 0.1% glutaraldehyde in PBST for 30 minutes at room temperature. The reverse transcription primer (‘oRT_CROPseq’, Key Resources Table) was synthesized at 100 nmol, purified by RNase Free HPLC, and supplied in IDT LabReady format (100 uM in IDTE, pH 8.0). Each well was washed twice with 0.5 mL PBST and once with 1X Reverse Transcription buffer in UltraPure water. The reverse transcription reaction mix was assembled on ice as previously reported.

300 uL of reverse transcription mix was supplied to each reaction well, and all wells not undergoing reverse transcription were left in PBST + 0.8 U/uL RiboLock used during the phenotype imaging. The plate was parafilmed and placed in a humidified tissue culture incubator for 3 hours. The plate was then transferred to a flat-top thermal cycler (ThermoFisher, 4484078) set to 42°C for 30 minutes and covered with a fitted piece of styrofoam to minimize loss to condensation on the plate lid. While the thermal cycler has a heated lid, loosely capping with styrofoam minimized any pressure on the plate that might compromise the structural integrity of the glass bottom. The plate was returned to room temperature, washed 5X with PBST, and post-fixed in freshly prepared 3% formaldehyde and 0.1% glutaraldehyde in PBST for 30 minutes at room temperature. After another 3X PBST washes, the plate was returned to the flat-top thermal cycler for 5 minutes at 37°C and 90 minutes at 45°C to complete padlock hybridization and gap-fill as previously reported25, scaled to a volume of 300 uL per well of a 12 well. The padlock probe (‘oPD_CROPseq’, Key Resources Table) was synthesized at 100 nmol, PAGE-purified, and supplied in IDT LabReady format. The sample was washed 3X with PBST and the final wash exchanged with 400 uL per well of rolling circle amplification mix, prepared as previously described25.

The parafilmed plate was returned to the flat-top thermal cycler for a 13 hour overnight incubation at 30°C. As during the gapfill reaction, a fitted piece of styrofoam was placed atop the plate to minimize volume loss to condensation on the lid. The next morning, wells were washed 3X with PBST prior to hybridizing the sequencing-by-synthesis primer at 1 uM in 400 uL 2X SSC per well for 30 minutes at 37°C on the flat-top thermal cycler. The SBS primer (‘oSBS_CROPseq’, Key Resources Table) was synthesized at 100 nmol, PAGE-purified, and supplied in IDT LabReady format (100 uM in IDTE, pH 8.0).

All washes in the following incorporation and cleavage steps were carried out using 600 uL room-temperature PR2 buffer per well. After aspirating the hybridization mixture, the first cycle was incorporated by supplying each well with 300 uL of freshly thawed and thoroughly inverted incorporation mix and incubating the plate for 3 minutes at 60°C. PR2 wash buffer was added to dilute the incorporation mix before aspiration, quickly followed by 6 additional PR2 washes at room temperature. The plate was washed an additional 5X with PR2 at 60°C for 5 minutes per wash.

The final P2R wash was replaced with 2X SSC with 200 ng/uL DAPI and incubated for 10 minutes to allow for saturation of the DAPI signal prior to imaging. Genotyping images were collected with channel-specific filters at 10X magnification (Key Resources Table) with the following run-specific settings: DAPI: 408 nm laser, 10% power, 15 ms exposure time; G and T: 546 nm laser, 30% power, 200 ms exposure time; A: 546 nm laser, 30% power, 200 ms exposure time; and C: 546 nm laser, 30% power, 350 ms exposure time.

2X SSC + DAPI imaging buffer in each well was exchanged for 300 uL of cleavage mix and incubated at 60°C for 6 minutes to cleave off the fluorescent terminator. Wells were washed quickly with PR2 3X at room temp, 3X at 60°C for 1 minute, and finally 3X at room temperature. Incorporation, imaging, and cleavage were repeated an additional 9 times for a total of 10 cycles.

### Single cell RNA sequencing

Cells were lifted with TrypLE Express (Thermo Fisher Scientific, 12605010), pelleted for 5 minutes at 300 × g, and washed in PBS with 0.04% BSA. Cells were diluted and strained through a 40 μm strainer. Cell viability was estimated to be around 90% per sample (Countess II, ThermoFisher). Prior to processing, a reference cell pool transduced with AAVS1-targeting guides (Table M1) was spiked into each sample such that 1000 singlets from the reference pool could be recovered per GEM well. These cells were used as an internal control to minimize batch effects between GEM wells during computational analysis.

**Table M1.**
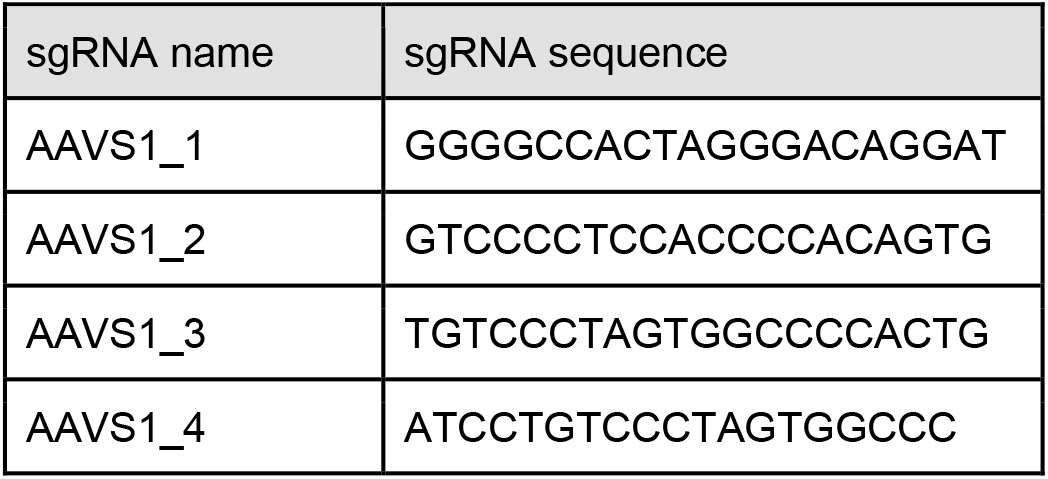
Reference cell pool sgRNA sequences.

Samples were processed with the Chromium Next GEM Single Cell 3’ HT Kit (10x Genomics, CG000416 Rev C). Pooled tag libraries treated with HyT36 were loaded across 3 GEM wells at 88 k cells each per timepoint with the goal of recovering 39 k singlets per well after filtering. Vehicle and untagged controls were loaded each into a single well at 30.5 k and 14.5 k cells, respectively, with the goal of recovering 17 k and 8 k singlets after filtering (Table M2).

**Table M2.** Single cell processing and sequencing parameters.

| # | sample | K cells loaded | K singlets target | M reads target |
| --- | --- | --- | --- | --- |
| 1 | TagLib_HyT36_2hr_1 | 88 | 39 | 1500 |
| 2 | TagLib_HyT36_2hr_2 | 88 | 39 | 1500 |
| 3 | TagLib_HyT36_2hr_3 | 88 | 39 | 1500 |
| 4 | TagLib_HyT36_12hr_1 | 88 | 39 | 1500 |
| 5 | TagLib_HyT36_12hr_2 | 88 | 39 | 1500 |
| 6 | TagLib_HyT36_12hr_3 | 88 | 39 | 1500 |
| 7 | TagLib_DMSO_2hr | 30.5 | 17 | 571 |
| 8 | UnTagLib_HyT36_2hr | 14.5 | 8 | 257 |
| 9 | targeted_sgRNA_libs | n/a | n/a | 165.8 |

Targeted amplification of sgRNA sequences from the CROPseq backbone (Addgene, 86708) was performed in order to reliably identify the tag in each cell. Sequencing libraries were prepared from three sequential PCR reactions starting from cDNA obtained during the 10x single cell processing protocol. The number of cycles for each PCR reaction was optimized by qPCR with EvaGreen (Biotium, 31000) on a CFX Opus 96 Real-Time PCR System (Bio-Rad).

• PCR #1: 10 μl reactions were prepared for each sample with KAPA HiFi HotStart ReadyMix (Roche, KK2602) containing 1 μl cDNA template and 1.5 μl of 10 μM PCR1 primer mix (U6 outer: TTTCCCATGATTCCTTCATATTTGC; TruSeq inner: CTACACGACGCTCTTCCGATCT). Cycling conditions: 95°C for 5 minutes, 17 cycles of 95°C for 30 seconds and 65°C for 45 seconds. PCR cleanup was performed with AMPure XP beads (Beckman Coulter, A63880) following manufacturer’s instructions.

• PCR #2: 10 μl reactions were prepared for each sample with KAPA HiFi HotStart ReadyMix containing 1 μl 25x diluted product from the previous step and 1.5 μl of 10 μM PCR2 primer mix (U6 inner: GTCTCGTGGGCTCGGAGATGTGTATAAGAGACAGCTTGTGGAAAGGACGAAACAC; TruSeq outer: ACACTCTTTCCCTACACGACGCTC). Cycling conditions: 95°C for 5 minutes, 13 cycles of 95°C for 30 seconds and 65°C for 45 seconds. PCR cleanup was performed with AMPure XP beads following manufacturer’s instructions.

• PCR #3: 10 μl reactions were prepared for each sample with KAPA HiFi HotStart ReadyMix containing 1 μl 25x diluted product from the previous step and 1.5 μl of 10 μM PCR3 primer mix (P7 and index: AATGATACGGCGACCACCGAGATCTACAC[10 bp Index]ACACTCTTTCCCTACACGACGCTC). Cycling conditions: 95°C for 5 minutes, 12 cycles of 95°C for 30 seconds and 72°C for 45 seconds. PCR cleanup was performed with 0.65x SPRIselect beads (Beckman Coulter, B23317) following manufacturer’s instructions. Products were quantified by Qubit 4 Fluorometer (Thermo Fisher Scientific) and pooled based on the ratio of singlets targeted (Table M2).

Transcriptome samples and the pooled targeted sample were mixed and sequenced on a NovaSeq 6000 with the 200 cycle kit (Illumina, 20028313). Final sample ratio was determined by the number of targeted reads as described in Table M2.

## QUANTIFICATION AND STATISTICAL ANALYSIS

### Tag library design and calculation of guide deviation score

The design and analysis of libraries for tagging is shown in Figure S1 with four primary steps: 1) selection of genes of interest, 2) identification of gene target sites, and 3) generation of lists of oligos. These steps have been condensed for broad and easy usage in the form of a genome-wide database of target sites at pooledtagging.org.

Genes were selected based on the specificity of their localization patterns and their level of expression. Specificity was determined based on available annotations on UniProt76. Expression was determined from RNA sequencing data acquired from the Human Protein Atlas77,78. Roughly 20 highly expressed genes were selected for each compartment. To identify all putative target sites available for intron tagging, the primary transcript for each gene was determined using the MANE database79 or APPRIS80 when a MANE entry was not available. Chromosomal exon coordinates for primary transcripts were acquired from Ensembl81 and converted into intron coordinates with coding sequence phase information. All intron coordinates were input into GuideScan82 to generate a list of all available target sites.

Up to 10 target sites were chosen per intron based on guide efficiency and specificity scores, and distribution throughout the intron. Distribution between the target sites was maximized using a recursive algorithm that prioritized top scoring guides. All target site sequences were divided into one of three groups based on intron phase, and negative control sgRNA sequences were added to each based on NTC guides from the Brunello CRISPR library73. To generate oligo tables, each sequence was concatenated with a restriction enzyme site and a unique complementary sequence for cloning as previously described83.

The guide deviation score depicted in Figure S1F-G is calculated in R as: mean(abs(*perfect_dist* - *diffs*)) / *perfect_dist*, where *diffs* is a vector of nucleotide spacing between each target in an intron, and *perfect_dist* = *intron_length* / (*target_count* - 1).

### Bulk sequencing of tagged library & feature analysis

sgRNA sequences were extracted from sequencing reads, aligned to a reference dataset of all possible sgRNAs, and demultiplexed using the CB2 package in R84. Samples of tagged pools after sorting were filtered to remove background sgRNA amplification; this was done by applying empirically-determined count thresholds that removed the majority of non-targeting control sgRNAs. Filtered samples were normalized using the formula: (reads per sgRNA / total reads per sample) × sample library size. Filtered and normalized phase-specific sublibraries were added together for analysis of tagging features.

Protein features such as relative surface accessibility, disorder, and structure were predicted for each residue using NetSurfP-2.085. Epigenetic marks were acquired from the corresponding hg38 tracks on the UCSC Genome Browser from K562 cells86, and all signals within 1 kb of the target site were summed. All other features, including sgRNA specificity and efficiency scores, protein lengths, and intron lengths were acquired in the process of generating oligo libraries for pooled tagging.

### CRISPR KO analysis

Knockout (KO) sgRNA libraries were aligned and demultiplexed as described above in the section: *Bulk sequencing of tagged library & feature analysis*. Background sgRNAs with low counts in all time points were filtered out. Non-targeting control guides were used to perform Loess normalization for all time points with the *affy* (v1.68.0) package in R. For gene-level analysis, KO-based depletion was calculated as log_2_(NCg + 1 / NCg + 1), where NCg are mean normalized sgRNA counts for gene *g* at either the early time point (ETP) or after 3 weeks of culture (3W). Genes with at least a two-fold depletion were considered “essential” for Figure 2C.

### Assigning cell genotypes using in situ sequencing

Briefly, raw imaging data were processed using the functions and snakemake workflow management infrastructure available on the GitHub repo for Optical Pooled screens36. The analysis was executed within a dedicated conda environment hosted on the high-performance computing resources at the Children’s Hospital of Philadelphia. Minor customizations to the underlying functions were made to calculate and report additional cell features and custom snakemake workflows were written to separately segment and extract features from 10X genotyping images and 40X phenotyping images (deposited on GitHub). Cell segmentation at both magnifications was carried out using CellPose’s ‘cyto’ model for the large-scale compartment-specific library and ‘cyto2’ for all subsequent optical sequencing runs87,88. The stepwise image processing workflow to call sgRNA reads from raw imaging data has been described extensively elsewhere25,36.

After read-calling, genotypes were mapped to phenotypes using a global coordinate framework to relate a plate-based location at 10X to the same location at 40X. This process and relevant software packages are described in detail in the Supplementary Text. In brief: the metadata for each image were scrubbed to collect the global, stage-based *x*,*y* position at the center of each image. The pixel-based coordinates reported for the centroid of each nucleus within the image were then used to assign each cell a stage-relative coordinate. Then, for each well, one cell in each of eight 10X tiles distributed evenly along the perimeter of the well was identified. The fiducial cell’s global coordinate in the 10X image was used as an anchor to identify the nearest-neighbor 40X tile, which was manually inspected to identify the corresponding cell. After establishing eight such ‘fiducial’ cells, the translation, rotation, and scaling transformations required to generate the 40X coordinate set from the 10X coordinate set were calculated by minimizing the least-square error between the algorithm-predicted and user-supplied 40X coordinates. The resulting mapping parameters were evaluated for accuracy on a randomly selected set of twenty cells; in the rare instances where the correct nucleus at 40X was not identified for every 10X cell, additional fiducial cells were added in increments of four and the parameters recalculated until complete accuracy was achieved.

Albums of cell images reflecting the localization patterns of a specific sgRNA were generated by printing out the area surrounding the mapped 40X nuclear coordinates with padding of 80 pixels in *x* and *y* directions (Fig. 7E; S8; Supplementary Text). Pre-calculated 40X cell masks and corresponding features were associated with 10X sgRNA reads by either 1) identifying the nuclear centroid at 40X that is the nearest neighbor to the 40X coordinate calculated using the mapping degrees of freedom or 2) intersecting the 40X nuclear masks with the calculated 40X coordinate. Jupyter notebooks for facile identification of fiducial cells and performing 10X to 40X mapping will be deposited on GitHub.

### Quantitation of localization patterns

To minimize the effects of genotyping errors on the quantification of localization patterns, we developed an unbiased approach to filter out cells that deviated from the most common localization for a given sgRNA. A convolutional neural network was employed to classify each cell image, bounded by its cytosolic segmentation mask, into one of five rough categories: ‘cellular diffuse’, ‘cytoplasmic features’, ‘nucleoplasm’, ‘nucleoli’, ‘nuclear membrane’. The network was trained on approximately 1000 manually curated cell images for each category (Supplementary Text). The modal category was considered representative of that sgRNA, and cells matching other patterns were removed from downstream analyses.

We used these classifier-filtered cells to train and employ a variational autoencoder with the goal of extracting latent parameters that capture localization patterns. To this end, we trained cytoself38 on a maximum of 1000 classifier-filtered cells per sgRNA of interest in the compact cell library (Supplementary Text). The trained cytoself network was then used to extract latent parameters from all classifier-filtered cells. These latent parameters were flattened and embedded onto a two-dimensional UMAP using UMAP-learn89, and UMAP clusters were identified and labeled using HDBSCAN40 with ‘min_cluster_size’ set to 800, ‘cluster_selection_epsilon’ set to 1.5.

### Processing of single cell RNA sequencing data

Gene expression and guide libraries were processed using the standard CellRanger (v6.1.1) workflow for single cell RNA sequencing (scRNAseq) with CRISPR libraries90. Briefly, reads were aligned to the GRCh38 assembly and protospacer calling for each barcode was determined via a Gaussian Mixture Model. Gene expression UMIs were quantified using UMI-tools (v1.1.4). Each GEM well was spiked with a cell population containing unique CRISPR guides meant to serve as a control. This population from each well was normalized using ComBat91 as implemented in the SVA package (v3.42.0) and the estimated hyperparameters were applied to the remaining cells per well. Putative doublets were identified using DoubletFinder92 (v2.0.3) and subsequent filtering was performed to obtain a count matrix retaining only cells with a single assigned sgRNA and normal mitochondrial (<12%) and ribosomal (>16%) RNA content. Principal component analysis of GEM well samples was performed using the R function *prcomp* on averaged gene counts per well.

### Tag group clustering and analysis

For tag group and gene comparisons, the data was processed as follows. sgRNAs were assigned to tag groups as described in the text. To calculate differentially expressed genes per sgRNA, specific cells were chosen from each timepoint (2 and 12 hours) to serve as a baseline control. We minimized tag-specific effects in control cells by sampling an approximately equal number of cells from each tag group followed by calculating a 0.5% trimmed mean. Gene fold changes per sgRNA were calculated by: (mean tag gene counts + 1) / (trimmed mean control gene counts +1). For each gene fold change, the Wilcoxon test *P* value was calculated using the *row_wilcoxon_twosample* function from the *matrixTests* (v0.1.9.1) package in R. In addition, for each sgRNA, the fold change and *P* value of the tagged gene were set to 1 to discount the effects of tagging on detection of transcripts. To calculate tag group-related fold changes, the mean gene fold changes across all sgRNAs per tag group were calculated and *P* values were combined using Fisher’s method via the *sumlog* function from the *metap* (v1.8) package in R. *P* values were adjusted for multiple comparisons using Hochberg’s method with the *p.adjust* R function, the minimum corrected *P* value for each gene across tag groups was determined, and all genes with a minimum corrected *P* value >0.001 were filtered out, leaving 3427 genes. Finally, to enable downstream comparison of genes and tag groups, log2 fold changes for each gene were scaled and centered across tag groups and timepoints with the *scale* R function. Centering was omitted in certain figures to avoid misinterpretation of fold change direction (Fig. 5C-E, Fig. S15A, Fig. 6, Fig. 6S).

Tag group-specific differentially expressed genes (Fig. S12) were determined using a two-sample Wilcoxon rank sum test comparing gene fold changes in a given tag group at both timepoints to fold changes in all other tag groups and timepoints with the *wilcox.test* R function. Genes with a *P* value < 0.05 were considered compartment-specific.

For Figure 5A, tag groups were compared by pairwise Pearson correlation with the *cor* R function. Groups were clustered with the *pheatmap* (v1.0.12) R package using the “average” clustering method. The calculated label order was used for all representations of the tag groups. For comparing tag group metagroups by gene set, as in Figures 5B and S13, each gene of a gene set was plotted in cartesian space with the fold change in metagroup 1 and 3 on the *x* and *y* coordinates, respectively. The significance of metagroup bias for each gene set was calculated with a one sample t test along the axis defined by the vector between origin and the gene set center, with *μ* = 0. Multiple comparison testing was performed with Hochberg’s method as described above, and the “metagroup difference score” was defined as -log_10_(FDR) for each gene set. The “tag group specificity score” was calculated as the ratio between the max and mean absolute fold change for each gene set to the fourth power.

### Two dimensional embedding of gene profiles

Minimum distortion embedding (MDE) and clustering of genes was performed as described previously55. To construct the embedding, *pymde* (v0.1.18) was initialized using the spectral embedding of the dataset (using sklearn’s SpectralEmbedding with n_components=19, affinity=’nearest_neighbors’, n_neighbors=34, eigen_solver=’arpack’)), and the *pymde* ‘‘preserve_neighbors’’ function was run with embedding_dim=2, n_neighbors=58, and repulsive_fraction=26. Following, clustering was performed using HDBSCAN with the parameters min_cluster_size=6, min_samples=2, cluster_selection_epsilon=0.2840.

Functional enrichment was performed on HDBSCAN gene clusters (Fig. 6A), compartment-specific gene sets (Fig. S12), and the top ∼100 upregulated genes for each sgRNA (Figs. 4D, S11). Significant GO:CC and GO:BP terms were determined using gProfiler93 with the *gprofiler2* (v0.2.1) R package using the parameters organism = “hsapiens”, correction_method = “g_SCS”, user_threshold = 0.08, ordered_query = F. For Figure S11, all GO:CC terms related to the row compartment were filtered and an average *P* value was calculated. Enrichment of protein quality control gene sets curated by the Proteostasis Consortium43,44 were determined using a hypergeometric test with the *dhyper* R function.

### Bulk RNA sequencing and correlations

For Bulk RNA sequencing experiments presented in Figure S4, data were processed using the nf-core/rnaseq pipeline94 (v3.9). Briefly, quality control was carried out using FastQC (v0.11.9). High quality reads were then aligned to the Ensembl GRCh38 reference genome index using both STAR95 (v2.7.10a) and Salmon96 (v1.2.0) for alignment and quantification. The following software versions were also used in the pipeline: Samtools97 (v1.15.1), RSeQC98 (v3.0.1), Qualimap99 (v2.2.2-dev), and Preseq100 (v3.1.1). Correlation between mean transcript counts and mean sgRNA tag counts per gene was calculated using the *cor* function in R using the Pearson method.

