## Supplementary Figures for "Pooled tagging and hydrophobic targeting of endogenous proteins for unbiased mapping of unfolded protein responses"

[7]Lead contact

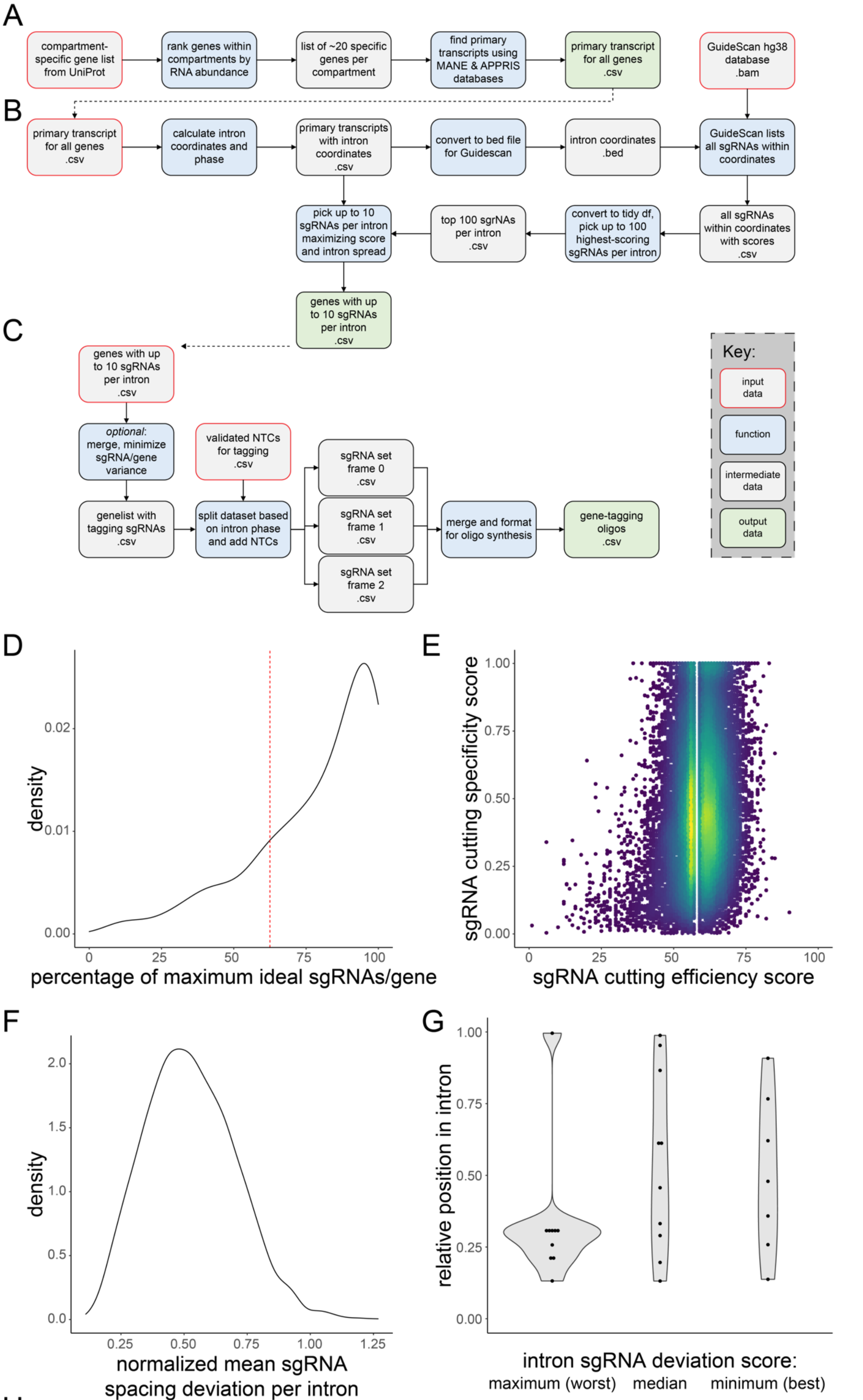


H

| # genes | # introns | # sgRNAs | avg ± sd sgRNAs per gene | % genes highly represented | mean efficiency score ± sd | mean specificity score ± sd | mean avg sgRNA deviation per intron ± sd |
|---|---|---|---|---|---|---|---|
| 265 | 2698 | 21345 | 80.5 ± 65.3 | 76 | 0.496 ± 0.264 | 58.9 ± 8.06 | 0.526 ± 0.178 |

**Figure S1. Design and analysis of oligo libraries for pooled tagging, related to Figure 1.**
A) Flowchart representing how targeted genes were selected. B) Flowchart representing how sgRNAs were picked for each gene. C) Flowchart representing how oligo libraries were designed from selected sgRNAs. D) The number of sgRNAs found per gene as a fraction of the ideal maximum number of sgRNAs, where each intron would have exactly 10 sgRNAs. E) Density plot of chosen sgRNA specificity and efficiency scores calculated by GuideScan. F) Metric representing how uniformly spread target sites are across a specific intron (STAR Methods). 0 means there is no deviation and guides divide an intron into segments of equal size. G) From F, sgRNAs mapped to introns with the maximum, median, and minimum deviation scores. H) Summary of metrics from the final sgRNA tagging library.

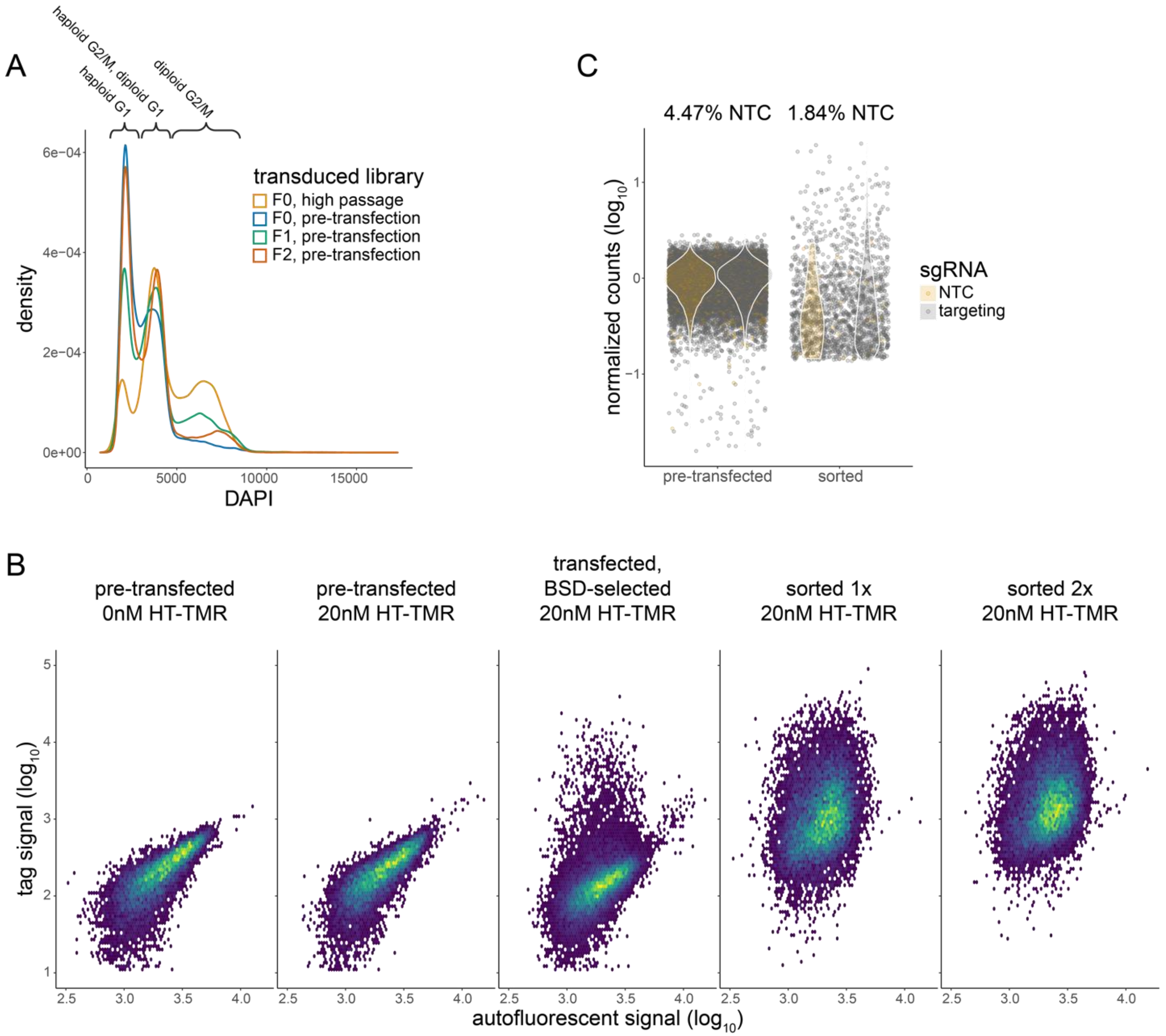


**Figure S2. Generation and tracking of pooled tag libraries in haploid HAP1 cells, related to Figure 1.**
A) DNA content analysis of transduced cells right before transfection. B) Flow cytometry of the F0 tag library at different stages of generation. C) Total count distributions of pre-transfected and sorted cells, labeled by targeting or non-targeting control (NTC) sgRNAs.

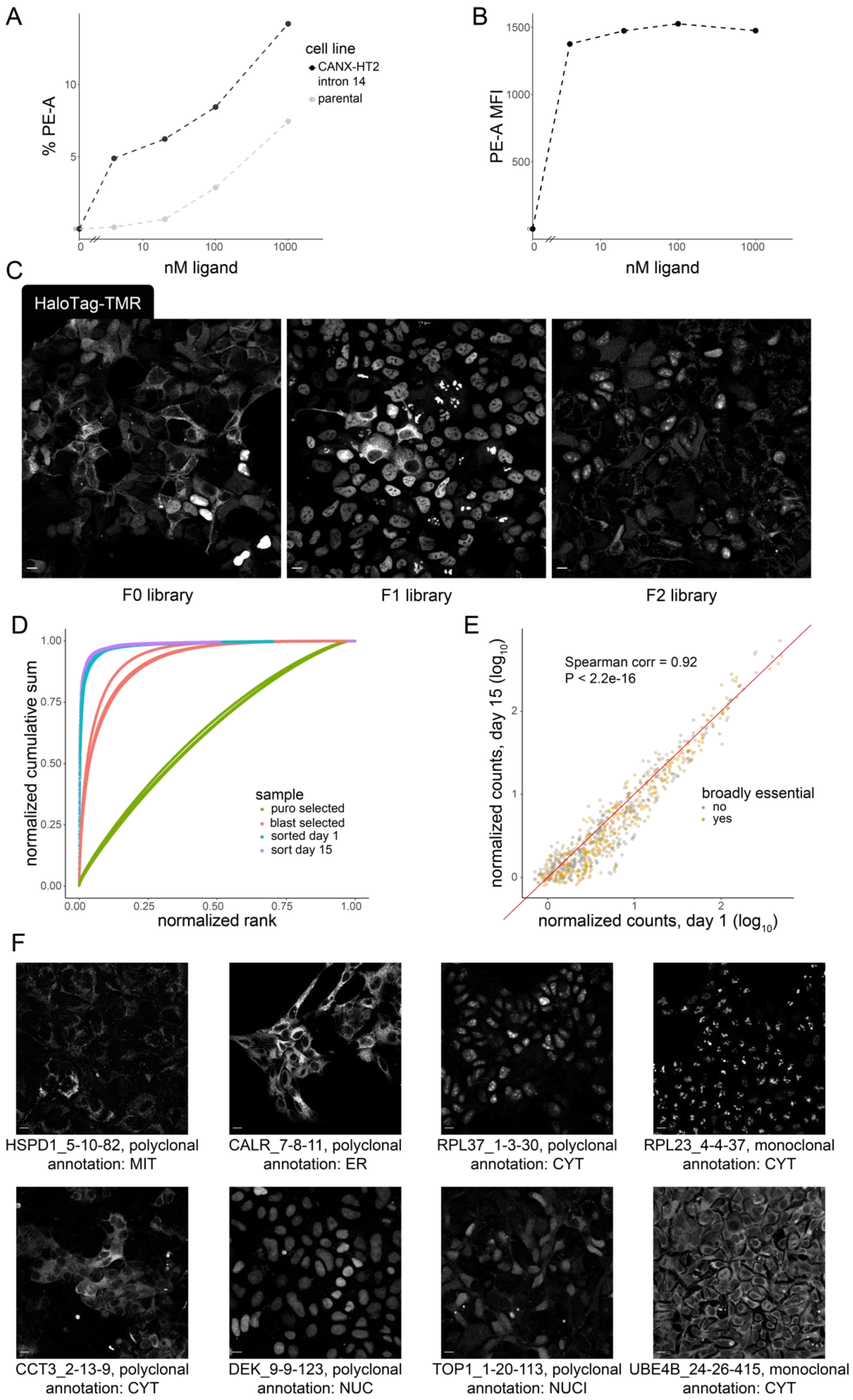
A
% PE-A
cell line
CANX-HT2 intron 14
parental
nM ligand
B
PE-A MFI
nM ligand
C
HaloTag-TMR
F0 library
F1 library
F2 library
D
normalized cumulative sum
normalized rank
sample
puro selected
blast selected
sorted day 1
sort day 15
E
Spearman corr = 0.92
P < 2.2e-16
broadly essential
no
yes
normalized counts, day 15 (log10)
normalized counts, day 1 (log10)
F
HSPD1_5-10-82, polyclonal
annotation: MIT
CALR_7-8-11, polyclonal
annotation: ER
RPL37_1-3-30, polyclonal
annotation: CYT
RPL23_4-4-37, monoclonal
annotation: CYT
CCT3_2-13-9, polyclonal
annotation: CYT
DEK_9-9-123, polyclonal
annotation: NUC
TOP1_1-20-113, polyclonal
annotation: NUCI
UBE4B_24-26-415, monoclonal
annotation: CYT

**Figure S3. Pooled tagging of endogenous proteins can be performed in HEK 293 cells and visualized with HaloTag-TMR, related to Figure 1.**
A-B) Dose curve of HaloTag-TMR treatment for 15 minutes before analysis by flow cytometry to assess A) the percentage of labeled cells gated in clonal populations of tagged or parental cells, and B) the mean fluorescence intensity (MFI) of the labeled sample. In B, non-specific labeling was filtered out by pulse-shape gating, demonstrating that tagged proteins are saturated with ligand even at low concentrations. C) Confocal images of pooled tag libraries in HEK 293 cells. D) Normalized cumulative sum and rank of all three HEK 293 libraries at different stages of generation. E) Comparison of sgRNA abundance in young and old total HEK 293 pooled tag libraries, colored by essentially of the tagged gene. F) Confocal images of HEK 293 cells generated by arrayed endogenous tagging using successful sgRNAs from the pooled tag libraries. The labels show the literature-annotated localization patterns and sgRNA names (*gene _ targeted intron # _ total # of introns _ sgRNA #).* Scale bars are 10 μm.

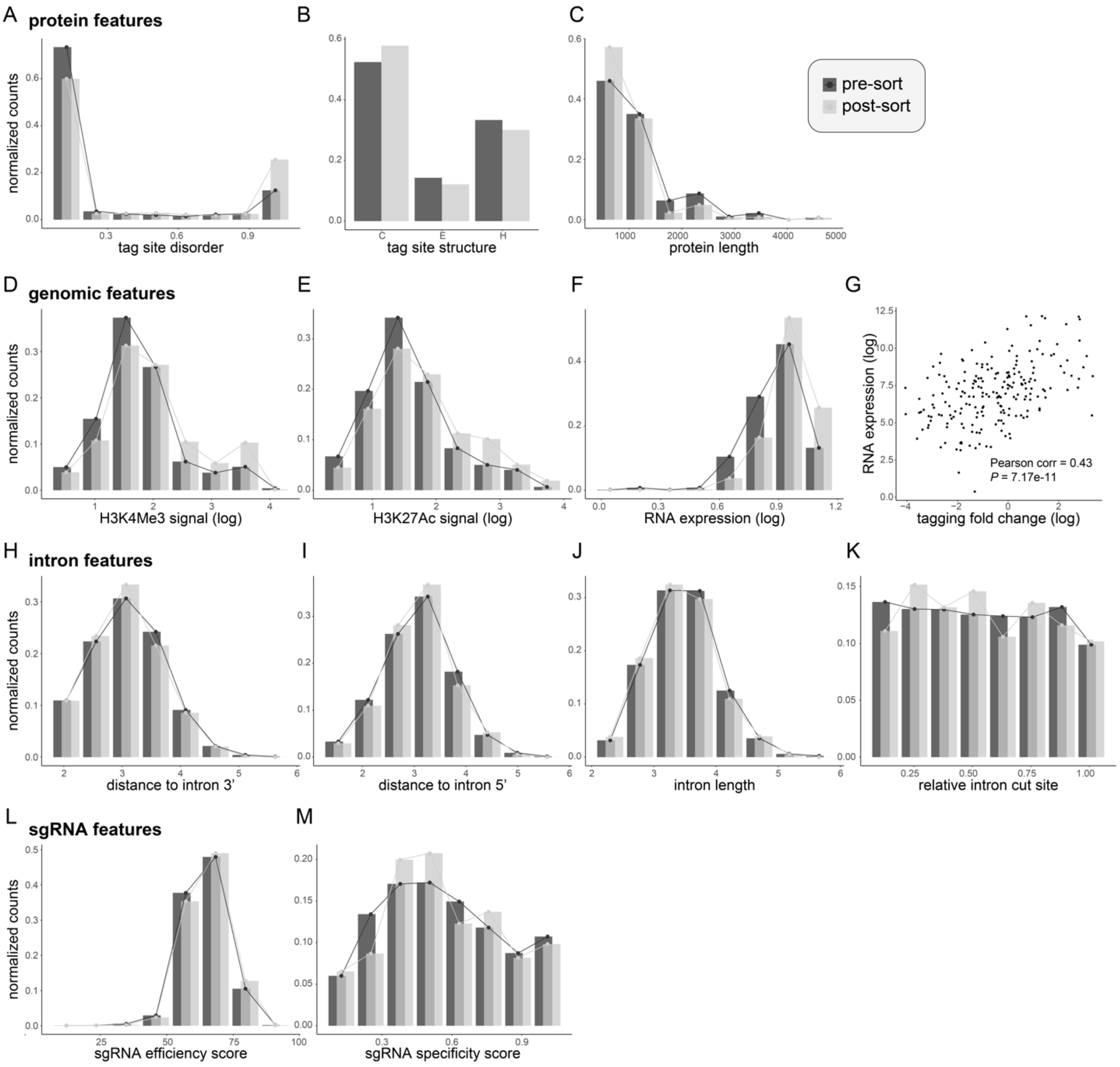


**Figure S4. Features that influence successful protein tagging, related to Figure 2.**
A-C) Effect of different protein features on the abundance of tagging sgRNAs, including A) local disorder, B) local secondary structure (C = coil, E = beta strand, H = alpha helix), and C) total protein length. D-E) Effect of different genomic features on the abundance of tagging sgRNAs, including D) target site H3K4Me3 signal and E) target site H3K27Ac signal. F-G) Effect of gene expression on F) sgRNA abundance and G) gene-level tagging fold change. H-K) Effect of different intron features on sgRNA abundance, including H) absolute distance to the 3' splice junction, I) absolute distance to the 5' splice junction, J) total intron length, and K) relative position of the target site within the intron. L-M) Abundance of sgRNAs with various cutting L) efficiency and M) specificity scores, pre- and post-sorting.

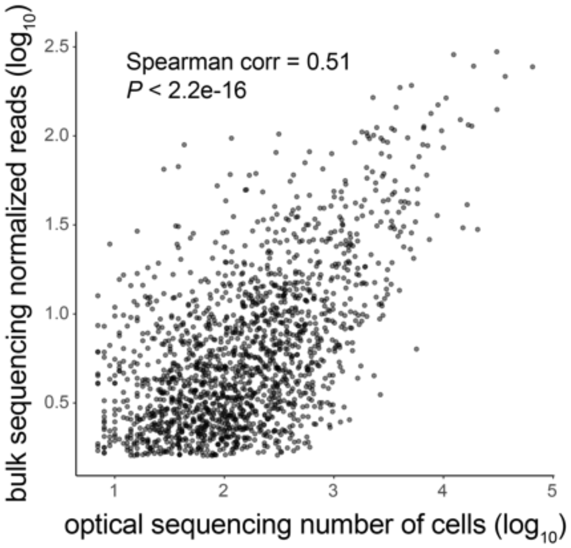


**Figure S5. Quantitation of sgRNA abundance in the pooled tagged library by optical sequencing and bulk NGS sequencing, related to Figure 3.**
The abundance of each sgRNA as measured by read counts from next-generation sequencing of sgRNA amplicons compared to the number of cells recovered for each sgRNA in the optical sequencing run.

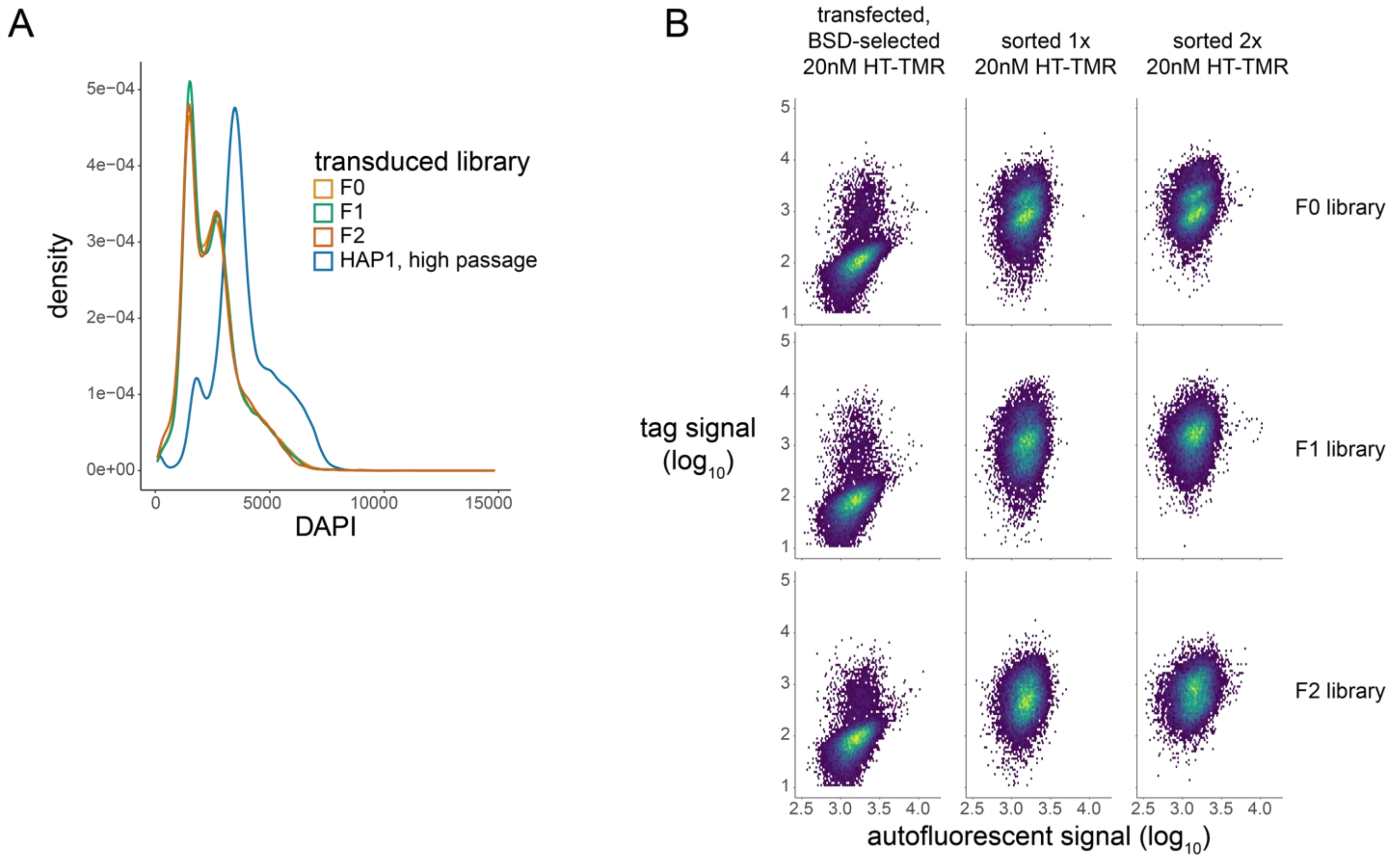


**Figure S6. Generation of the compact tag library in HAP1 cells, related to Figure 4.**
A) DNA content analysis of transduced cells right before transfection. B) Flow cytometry of the three phase-specific subsets of the compact tag library at different stages of generation.

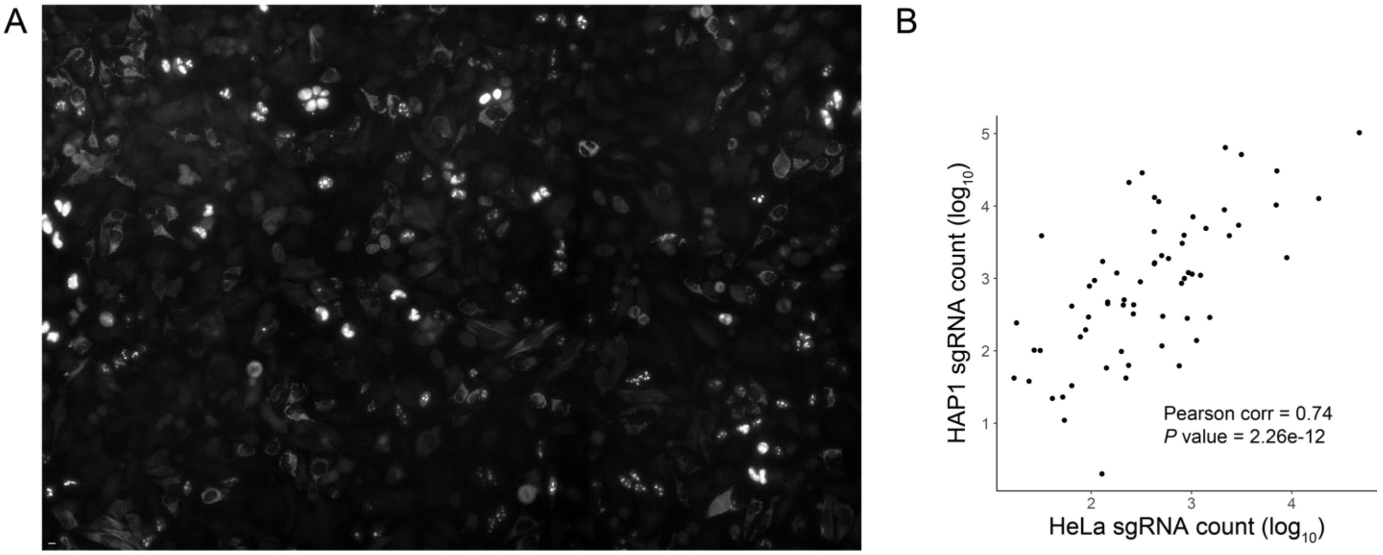


**Figure S7. Pooled tagging in HeLa cells performs similarly to that in HAP1 cells, related to Figure 4.**
A) Widefield fluorescence image of the compact F0 library generated in HeLa cells. Scale bar in the lower left represents 10 µm. B) Comparison of sgRNA abundance of the compact F0 library generated in HeLa versus HAP1 cells.

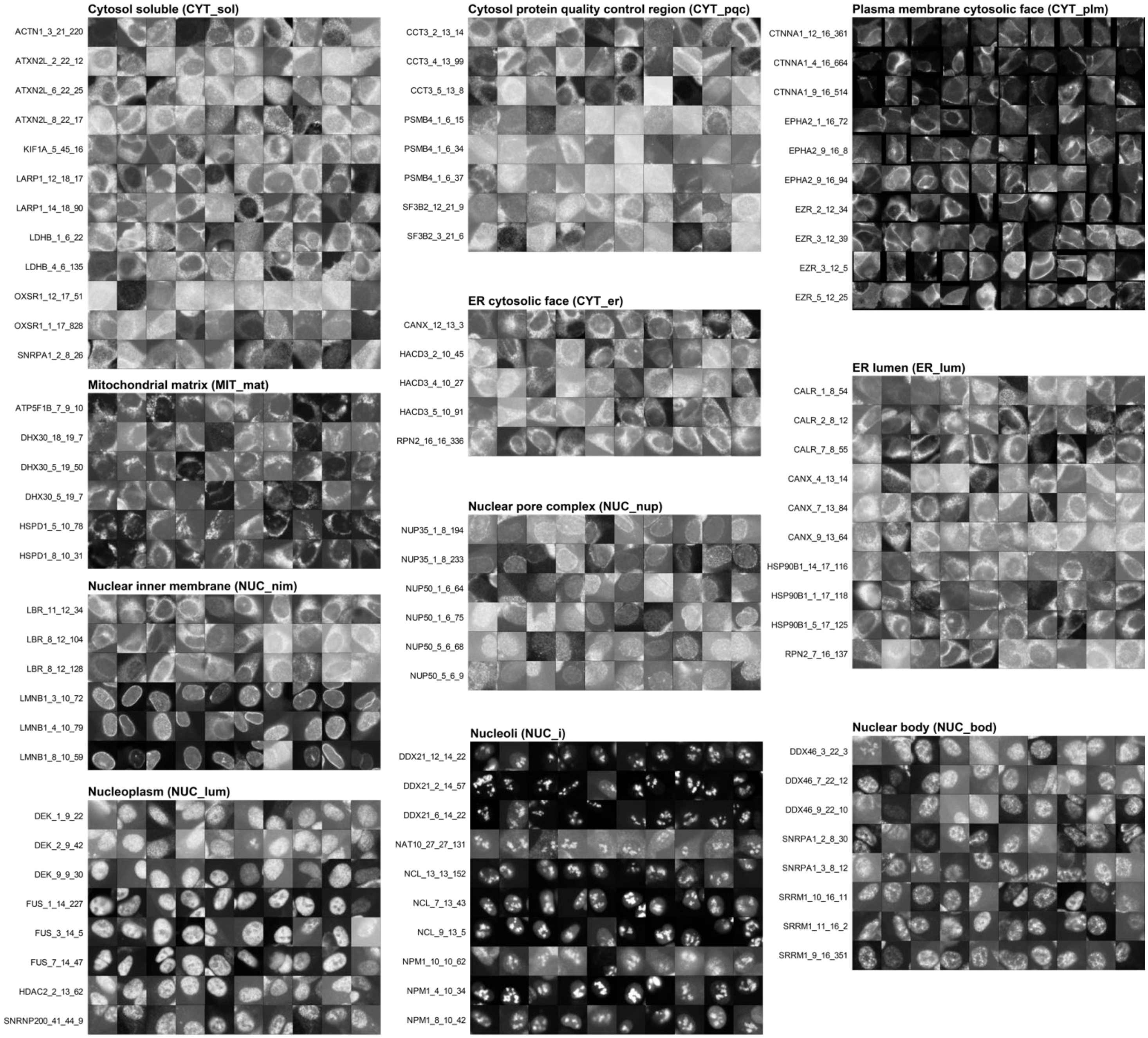


**Figure S8. Cell albums for each sgRNA from the compact tag library, related to Figure 4.**
sgRNAs are divided into compartment-specific tag groups and 10 cells are sampled for each.

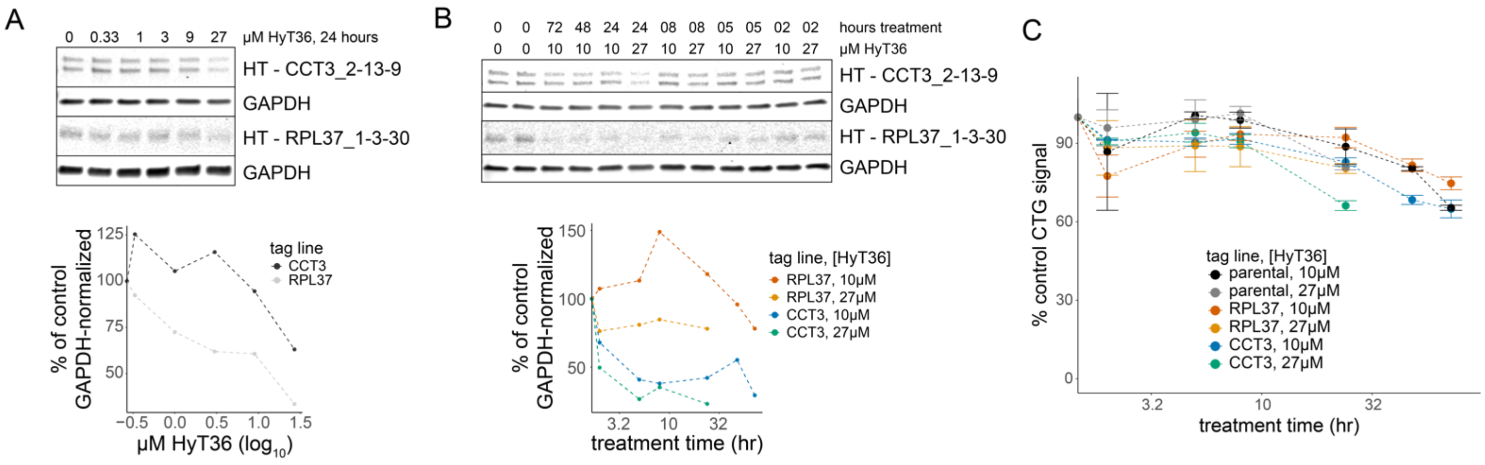


**Figure S9. Optimization of HyT36 for destabilization of endogenously tagged proteins, related to Figure 4.**
A-B) Immunoblot assay and quantitative analysis of clonal lines of tagged CCT3 and RPL37 treated with HyT36 at multiple A) concentrations and B) time points. C) Cell viability of parental and clonal tagged lines treated with HyT36 for up to 24 hours at 27 µM or 72 hours at 10 µM.

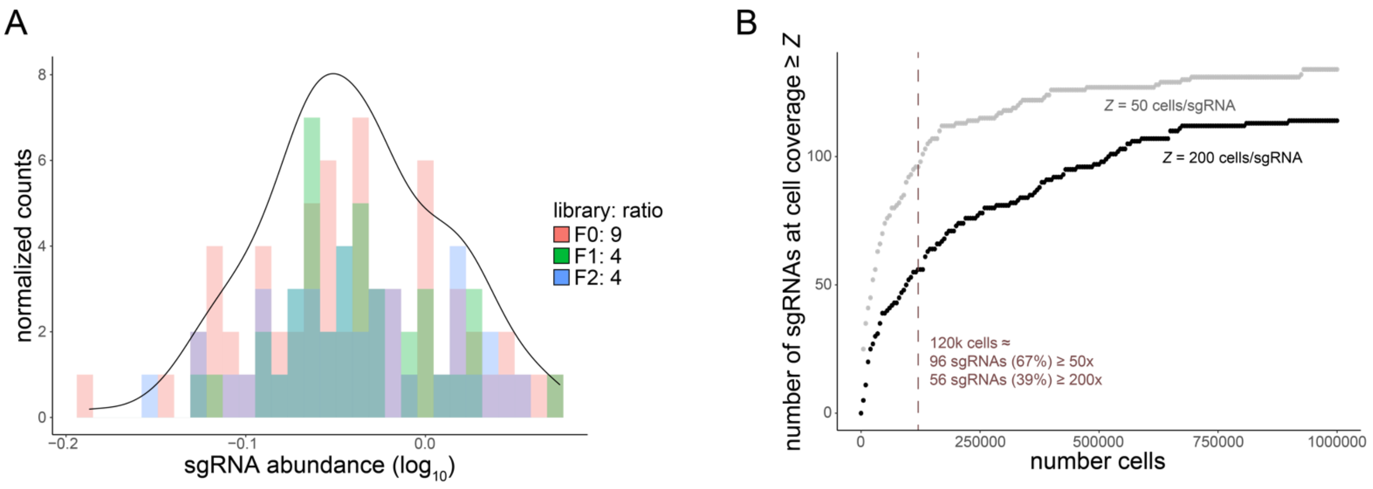


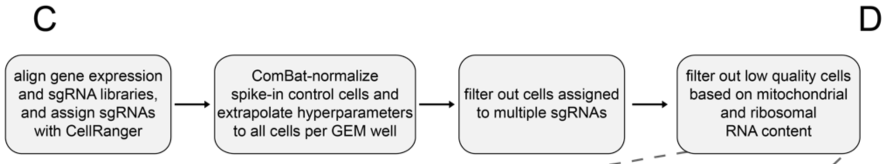


| sample | cells | number of tags | counts/cell | genes/cell |
|---|---|---|---|---|
| 2hr HyT36 well 1 | 25300 | 123 | 15403 | 4316 |
| 2hr HyT36 well 2 | 25368 | 120 | 14584 | 4160 |
| 2hr HyT36 well 3 | 22892 | 119 | 14933 | 4238 |
| 12hr HyT36 well 1 | 31140 | 126 | 14228 | 4146 |
| 12hr HyT36 well 2 | 32245 | 123 | 14291 | 4216 |
| 12hr HyT36 well 3 | 32100 | 125 | 12780 | 3934 |
| 2hr DMSO | 15153 | 110 | 10420 | 3547 |
| 2hr untag HyT36 | 7134 | 85 | 11360 | 3622 |

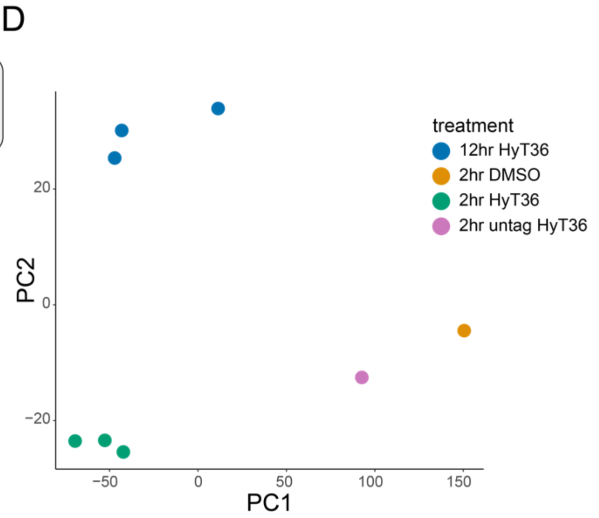


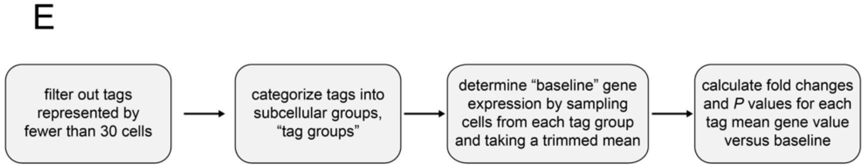


**Figure S10. Design and metrics for single cell RNA sequencing of destabilized SPOTLITES libraries, related to Figure 4.**
A) sgRNA distribution in phase-specific compact tag libraries combined at the indicated ratio. The outline represents the density distribution of the combined library. B) Required size of the combined compact tag library to achieve at least 50x (gray line) or 200x (black line) cell coverage for the indicated number of guides. C) Workflow for processing sequencing reads to a count matrix, and table describing the cells and tags recovered. D) Principal component plot to show similarity of GEM wells. E) Workflow to calculate tag-associated gene fold changes induced by HyT36 treatment.

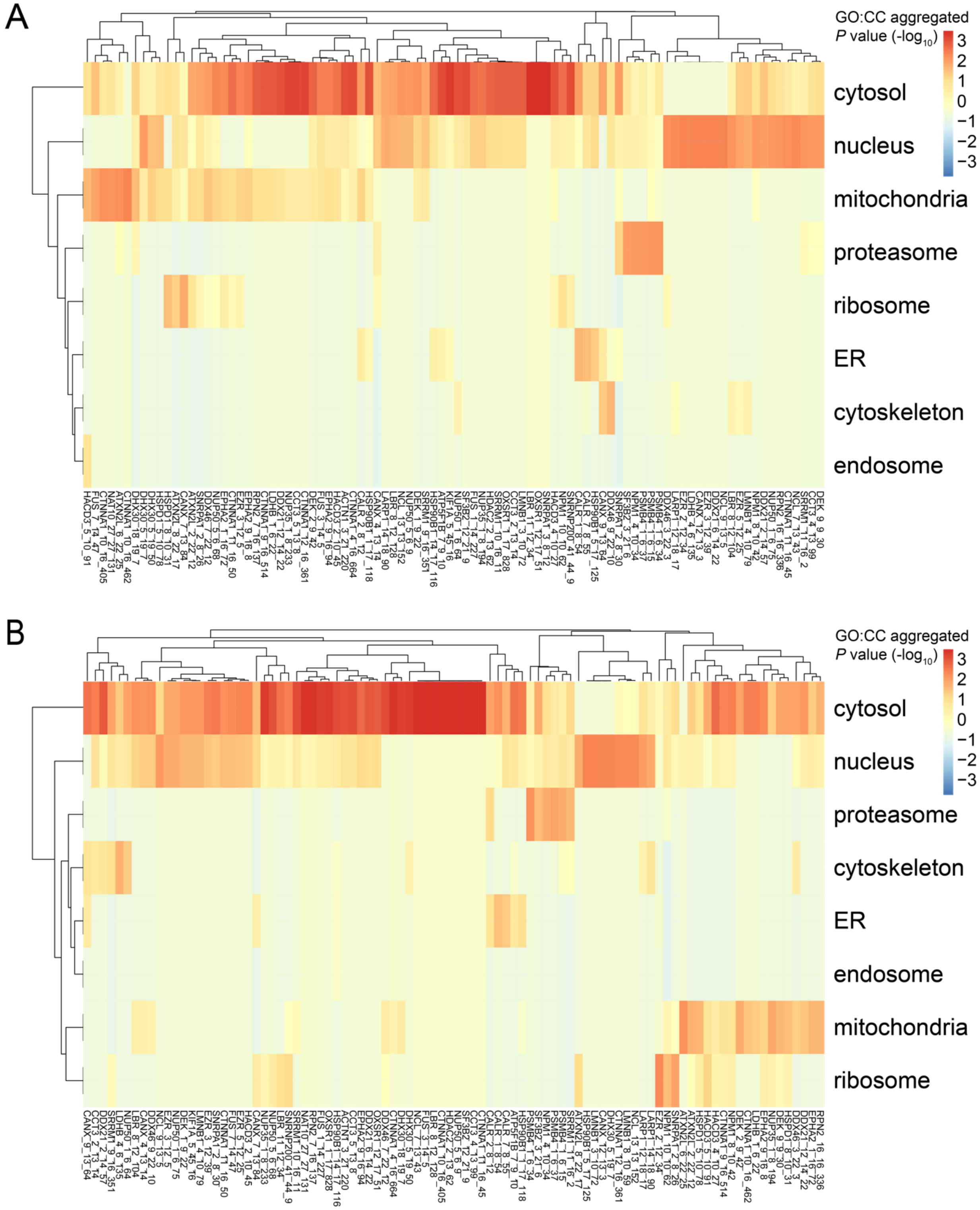


**Figure S11. Destabilization of compartment-specific tags tends to upregulate genes associated with the corresponding compartment, related to Figure 4.**

A-B) Heatmap of compartment-associated terms enriched within the top ~100 upregulated genes for each tag at A) 2 hours and B) 12 hours of HyT36 treatment. Values represent the average P value of all GO:CC terms related to the indicated compartment. sgRNA naming structure follows: *gene* _ *targeted intron #* _ *total # of introns* _ *sgRNA #*.

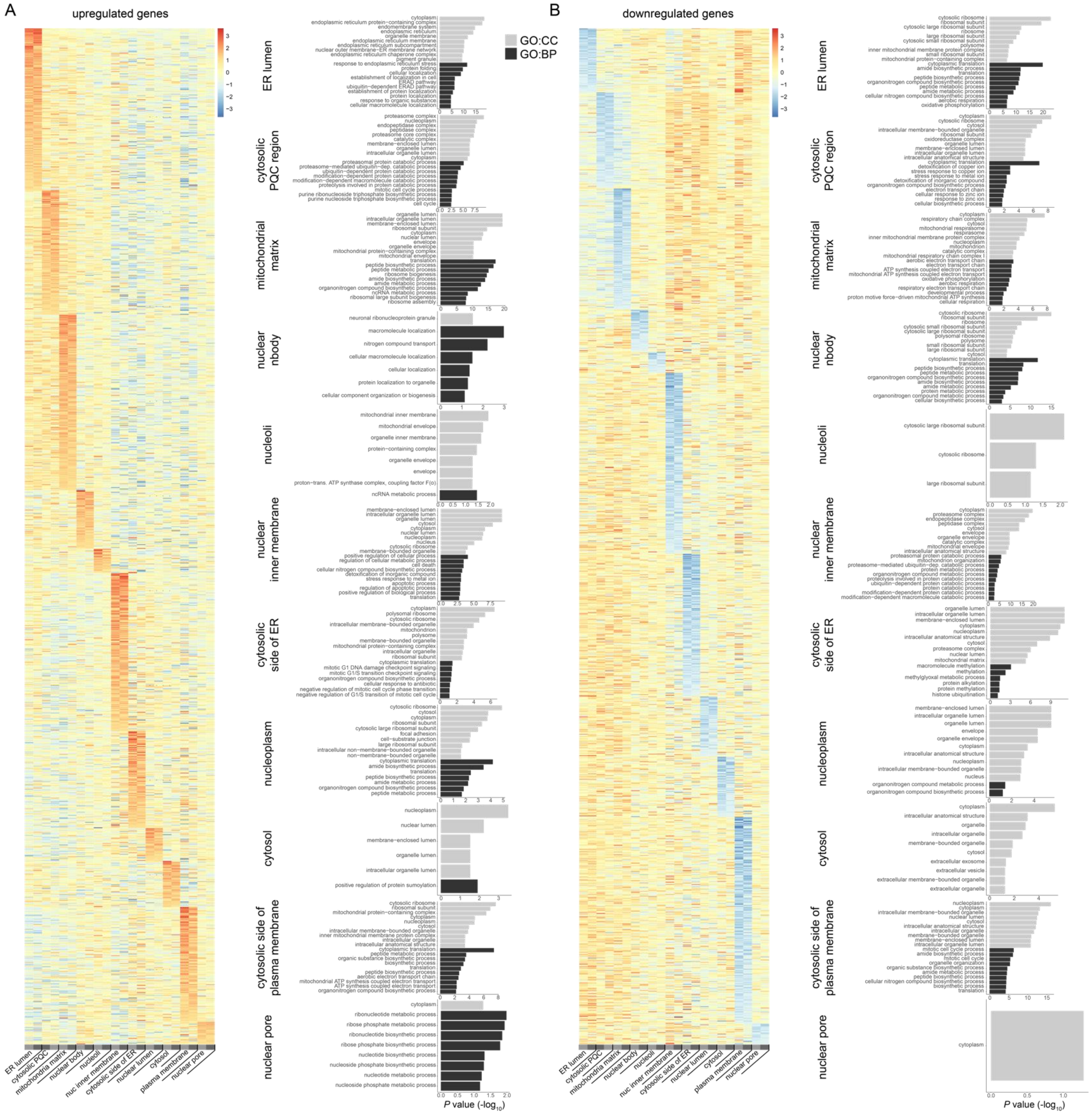

**Figure S12. Compartment-specific differentially expressed genes often have compartment-relevant functions, related to Figure 5.**

A-B) Heatmap of Z-normalized log fold changes across tag groups for compartment-specific A) upregulated and B) downregulated genes. Functional enrichment of GO:CC and GO:BP terms is performed for each compartment-specific gene set.

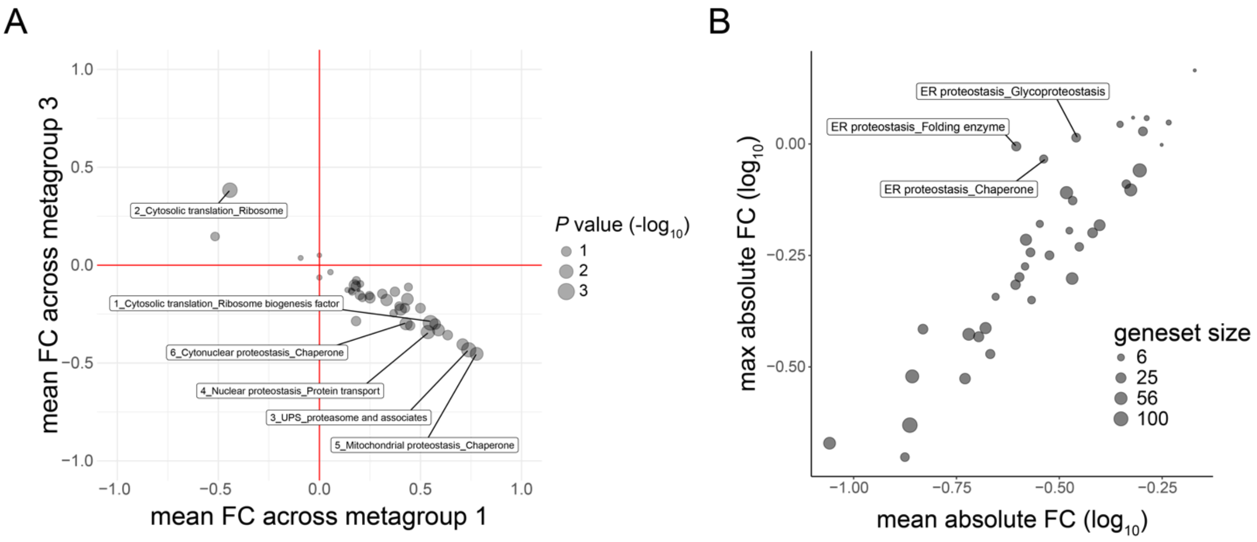


**Figure S13. Metagroups differ by expression of proteostasis-related genes, related to Figure 5.**
A) Scatter plot of average gene fold change values per proteostasis-specific gene set between metagroups 1 and 3. The P value indicates the significance of gene set differential expression between metagroups. B) Scatter plot of the mean versus maximum absolute fold change of a gene set across tag groups.

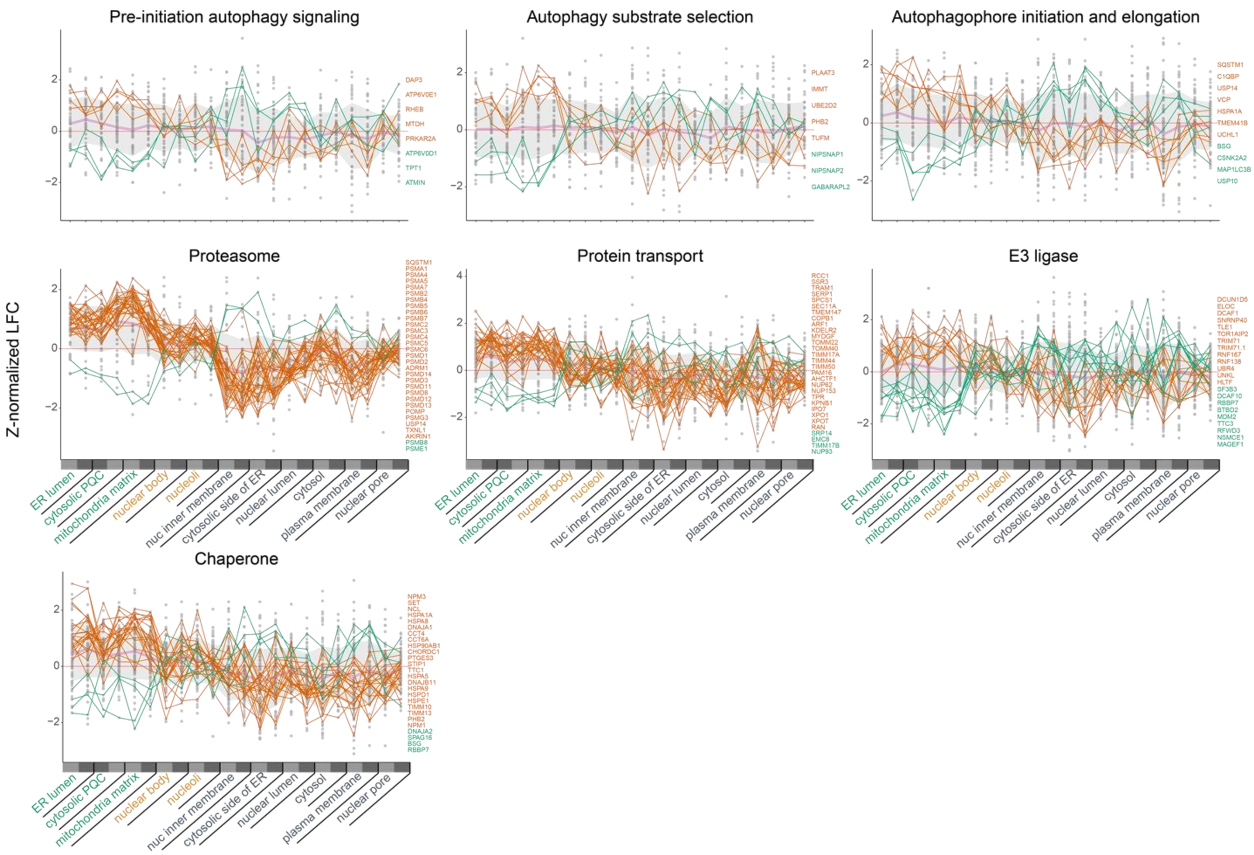


**Figure S14. Certain gene sets have gene subsets with inverse regulation patterns, related to Figure 5.**
Points represent each gene in the labeled gene set, the thick pink line represents the mean of points, and the gray ribbon represents one standard deviation from the mean. Genes with differential expression between metagroups 1 and 3 are labeled.

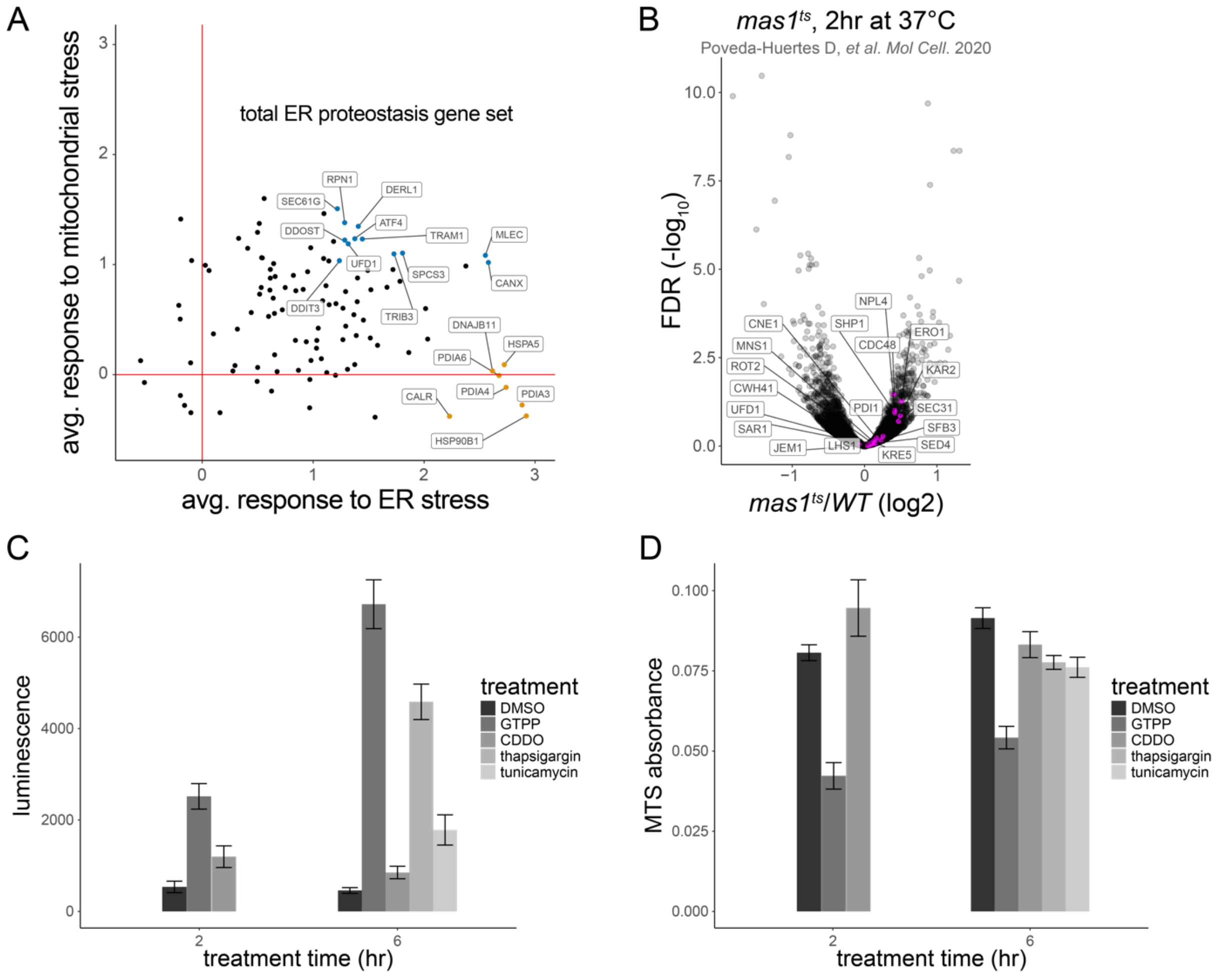


**Figure S15. Protein misfolding in the mitochondria directly induces the IRE1α branch of the ER UPR, related to Figure 5.**
A) Scatter plot of the average normalized fold change of all ER proteostasis genes between 2 and 12 hours of hydrophobic tagging. Highly upregulated genes exclusive to destabilization in the ER are colored orange, while shared genes are colored blue. B) Volcano plots of RNA sequencing data from the indicated study in yeast cells, with ER proteostasis-associated genes colored in magenta. C) MTS-based cell viability assay in response to treatments. D) Raw luminescence signal from XBP1 splicing luciferase reporter.

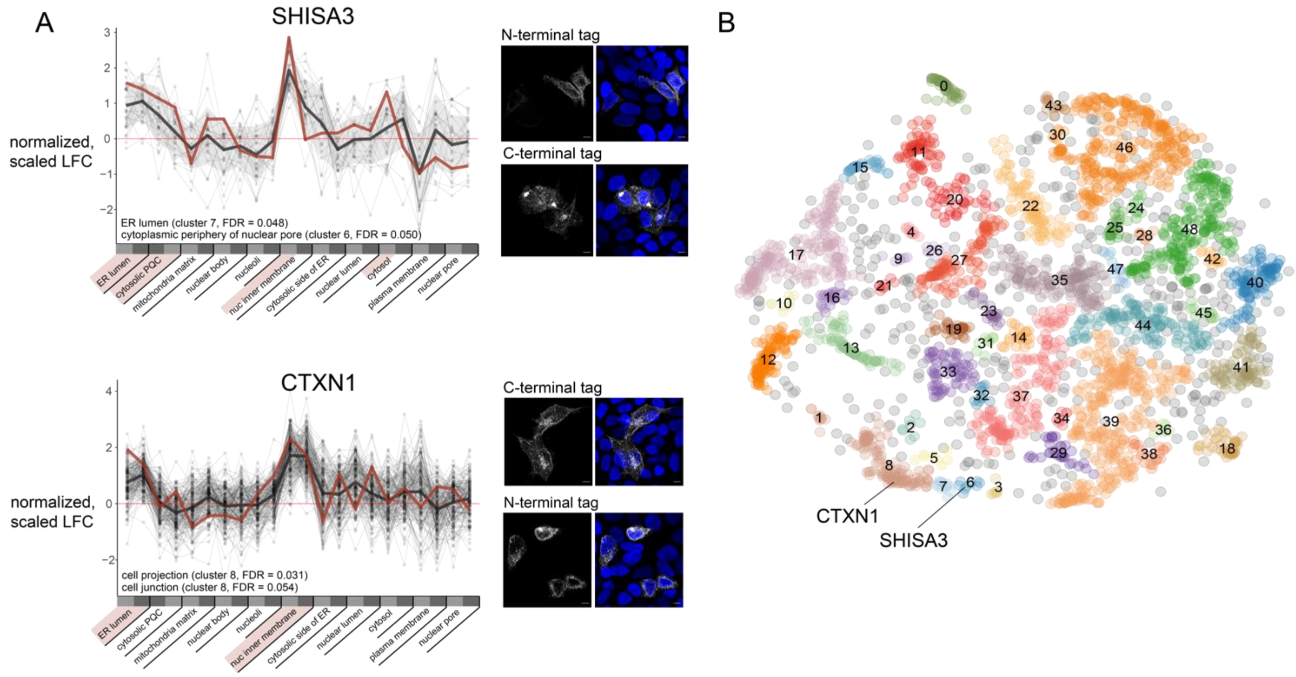


**Figure S16. Characterization of additional poorly annotated genes, related to Figure 6.**
A) Poorly annotated genes that were not candidates for SPOTLIES tagging were expressed in HAP1 cells with C- and N-terminal mClover3 fusions to determine localization patterns. The expression pattern of each gene in response to hydrophobic tagging across compartments is displayed to the left, with the fold-change of the unannotated gene in red, with all genes in the associated clusters in thin black lines and the average of those genes represented by a thick black line. In the case of SHISA3, small adjacent clusters 3, 6, and 7 were considered in aggregate, as SHISA3's cluster alone had too few genes for comprehensive recovery of functionally-enriched terms. Relevant functional enrichment terms associated with the clusters of each depicted gene are listed in the lower left-hand corner of the transcriptional profile plots. Confocal images of each tagged protein are overlayed with DAPI, with scale bars representing 10 μm. B) Minimum distortion embedding as in Figure 6A, highlighting the cluster location of genes in S16A.

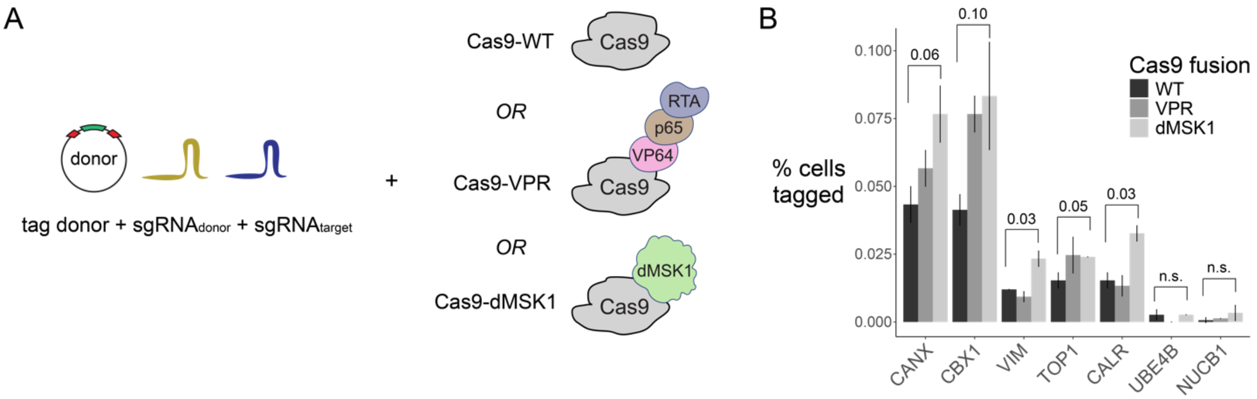


**Figure S17. Characterization of epigenetic activators, related to Figure 7.**
A) Cas9 fusions to several activator domains nominates B) dMSK1-Cas9 as a locus-generic reagent for increasing tagging efficiency.
