## Supplementary Text for "Pooled tagging and hydrophobic targeting of endogenous proteins for unbiased mapping of unfolded protein responses"

Sections:

1. Cross-objective mapping
2. Training and architecture of the convolutional neural network classifier
3. Training the variational autoencoder cytoself for unbiased clustering of cell images

### 1. Cross-objective mapping

*Scrubbing metadata from images*

Metadata detailing the x and y global stage position per tile in microns were scrubbed from raw imaging tifs (*tifffile* 2021.8.30) or from raw nd2 file (*pims-nd2* 1.1). Additional metadata collected include the pixel size in microns, which was used to calculate each segmented cell's global position in microns using tile-relative pixel coordinates. File labels were parsed for the tile number, which reflects which tiles are adjacent to one another; this information is later used to arrange the tiles on the global coordinate system.

*Identifying fiducial cells*

The approximate diameter of each well is calculated by subtracting the global micron y coordinate of the top-most tile from that of the bottom-most tile, and the centroid of the well is approximated by the mean of these two coordinates (*pandas* 1.0.3). The radius is calculated from the diameter. Candidate tiles in which to look for fiducial cells are identified by finding the tiles with centers closest to the following points, each of which is at 90% of radius of the well (*pandas* 1.0.3):

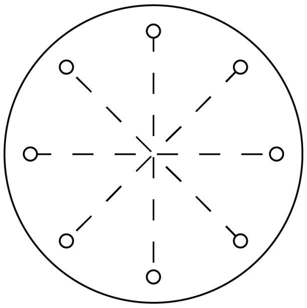

After identifying these fiducial tiles at 40X, the cognate 10X tile is identified by searching for the 10X tile center coordinates for the nearest neighbor to the center coordinates of the 40X tiles (*numpy* 1.18.1, *scipy* 1.6.0). Images of the 10X and 40X tile pairs are printed out in a jupyter notebook annotated with their cell IDs at each magnification, to allow the user to look up cell coordinates for a given cell ID (jupyter *notebook* 6.4.3, *numpy* 1.18.1, *matplotlib* 3.1.3, *pillow* 7.1.2). Thus coordinates are found for one fiducial cell per tile at 10X and at 40X.

*Arranging relative position of images on global coordinate map*

The tile number and center coordinates scrubbed from the filename and metadata above are used to arrange each tile in a square matrix representing the entire well (*pandas* 1.0.3, *numpy* 1.18.1). Empty positions in the square matrix where no tile exists (due to the circular well shape) are populated with -1. Neighboring tiles in the matrix are printed to a jupyter notebook as before and manually evaluated to determine how they must be oriented relative to one another in order to form a contiguous image.

*Calculating affine transformation matrix that minimizes distance between user-supplied fiducial cell coordinates and predicted fiducial cell coordinates*

Given the coordinates at both magnifications of the fiducial cells, five mapping parameters (x-translation, y-translation, angle of rotation, x-scaling, and y-scaling) for a given well are calculated using a sequential least square programming minimization function (*scipy* 1.6.0), and summarized in an affine transformation matrix (*numpy* 1.18.1). Once mapping parameters are optimized, the affine transformation is performed for every cell in the well, such that every segmented cell at 10X is ultimately associated with a tile number and pixel-based coordinate at 40X (*numpy* 1.18.1). These coordinates can be synchronized with information extracted from each image using the phenotyping and genotyping snakemake workflows (*snakemake* 5.32.1, our GitHub Repo).

*Evaluating accuracy of the transformation matrix using a randomly sampled set of cells*

A set of 20 randomly selected cells at 10X and their calculated 40X coordinates are displayed as described above. If the mapping has not successfully identified the center of the correct cell nucleus for all cells, additional fiducial cells may be added and the mapping parameters re-calculated.

**2. Training and architecture of the convolutional neural network classifier**

Training was performed using three channels: the HaloTag-fluor channel cropped to the cellular mask, the HaloTag-fluor channel cropped to the nuclear mask, and the HaloTag-fluor channel within the cellular mask after setting all pixels within the nuclear mask to 0. Manually curated cell images were assigned to training, validation, and test sets at proportions of 0.64, 0.16, and 0.2, respectively. Due to the small dataset and simplicity of the CNN, training time was on the order of several minutes using a Tesla P100 16 GB GPU, for 200 epochs, with a batch size of 256. The classifier architecture is as follows:

| Input | Layer | Activation | Pool | Output |
|---|---|---|---|---|
| 3 x 128 x 128 | Conv 32 x 3 x 3<br>Stride = 1<br>Padding =1 | ReLu | Filter = 3 x 3<br>Stride = 2 | 32 x 64 x 64 |
| 32 x 64 x 64 | Conv 64 x 3 x 3<br>Stride = 1<br>Padding =1 | ReLu | Filter = 3 x 3<br>Stride = 2 | 64 x 32 x 32 |
| 64 x 32 x 32 | Conv 128 x 3 x 3<br>Stride = 1<br>Padding =1 | ReLu | Filter = 3 x 3<br>Stride = 2 | 128 x 16 x 16 |
| 128 x 16 x 16 | Conv 256 x 3 x 3<br>Stride = 1<br>Padding =1 | ReLu | Filter = 3 x 3<br>Stride = 2 | 256 x 8 x 8 |

| | | | | |
|---|---|---|---|---|
| 256 x 8 x 8 | Conv 512 x 3 x 3<br>Stride = 1<br>Padding =1 | ReLu | Filter = 3 x 3<br>Stride = 2 | 512 x 4 x4 |
| 512 x 4 x4 | Flatten | - | - | 8192 |
| 8192 | FC | ReLu | - | 1024 |
| 1024 | FC | Softmax | - | 5 |

The following confusion matrix and metrics quantify the performance of the trained CNN:

Accuracy = 0.929, Precision = 0.931, Recall = 0.929, F1 = 0.930

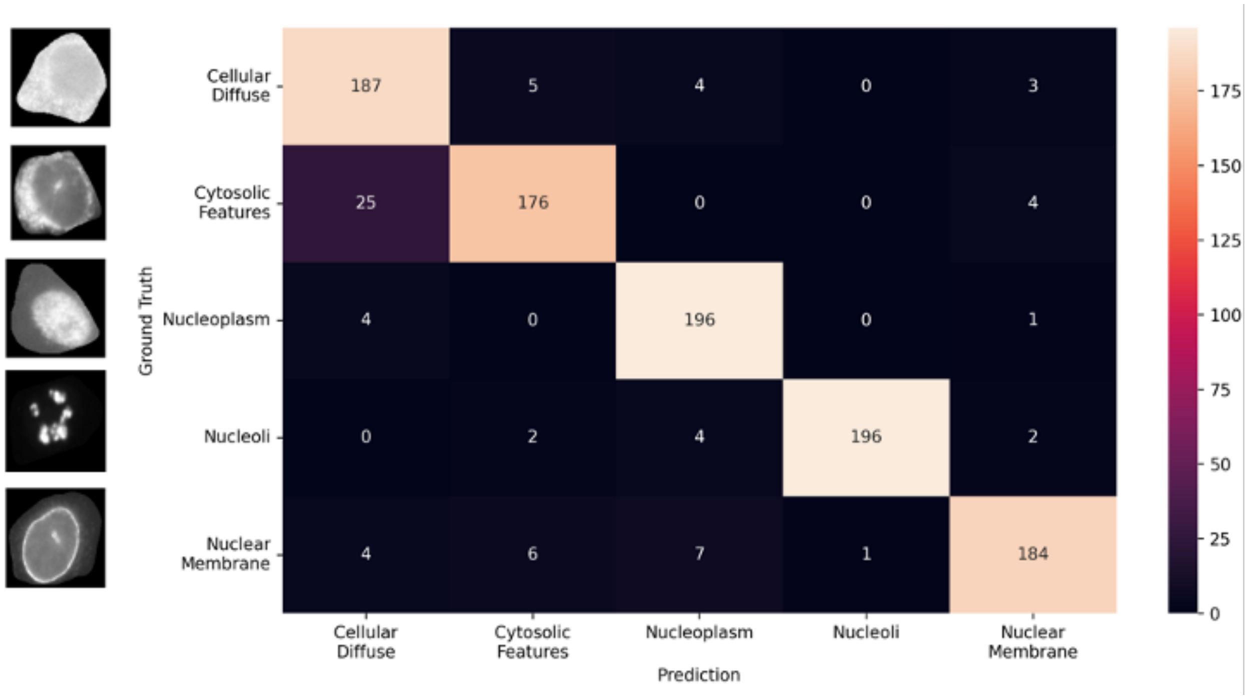


**3. Training the variational autoencoder cytoself**

Training was performed using the same three channels described in '2. Training and architecture of the convolutional neural network classifier.' Manually curated cell images were assigned to training, validation, and test sets at proportions of 0.7, 0.15, and 0.15, respectively. Cropped cell png images were resized to 128 x 128 pixels (*pillow* 7.1.2). Training was done on a A100-SXM4-40GB GPU with a batch size of 256. The training parameters were as follows:

| Parameter | Value |
|---|---|
| Input Shape | 3 x 128 x 128 |
| Embedding Shape | (32, 32), (4, 4) |

| | |
|---|---|
| Embedding Shape | 32 |
| Embedding Number | 512 |
| FC Output Index | 2 |
| Class Number | 12 |
| FC Input Tube | vqindhist |
| Learning rate | 1e-4 |
| Max Epoch | 100 |
| Reduce Learning Rate Patience | 3 |
| Reduce Learning Rate Increment | 0.1 |
| Early Stop Patience | 10 |
